# Mechanical comparison of *Escherichia coli* biofilms with altered matrix composition: a study combining shear-rheology and microindentation

**DOI:** 10.1101/2025.02.07.637005

**Authors:** Macarena Siri, Adrien Sarlet, Ricardo Ziege, Laura Zorzetto, Carolina Sotelo Guzman, Shahrouz Amini, Regine Hengge, Kerstin G. Blank, Cécile M. Bidan

## Abstract

The mechanical properties of bacterial biofilms depend on the composition and micro-structure of their extracellular matrix (ECM), which constitutes a network of extracellular proteins and polysaccharide fibers. In particular, *E. coli* macrocolony biofilms were suggested to present tissue-like elasticity due to a dense fiber network consisting of amyloid curli and phosphoethanolamine-modified cellulose (pEtN-cellulose). To understand the contribution of these two main ECM components to the emergent mechanical properties of *E. coli* biofilms, we performed shear-rheology and microindentation experiments on biofilms grown from *E. coli* strains that produce different ECM. We measured that biofilms containing curli fibers are stiffer in compression than curli-deficient biofilms. We further quantitatively demonstrate the crucial contribution of pEtN-cellulose, and especially of the pEtN modification, to the stiffness and structural stability of biofilms when associated with curli fibers. To compare the differences observed between the two methods, we also investigated how the structure and mechanical properties of biofilms with different ECM compositions are affected by the sample preparation method used for shear-rheology. We found that biofilm homogenization, used prior to shear-rheology, destroys the macroscale structure of the biofilm while the microscopic ECM architecture may remain intact. The resulting changes in biofilm mechanical properties highlight the respective advantages and limitations of the two complementary mechanical characterization techniques in the context of biofilm research. As such, our work does not only describe the role of the ECM on the mechanical properties of *E. coli* biofilms. It also informs the biofilm community on considering sample preparation when interpreting mechanical data of biofilm-based materials.

## Introduction

Biofilms are complex, highly heterogeneous living materials, composed of bacteria embedded in a self-produced extracellular matrix (ECM) of proteins and polysaccharides. In recent decades, biofilm mechanics has been studied across diverse fields, from microbiology to materials science and soft matter physics.^1^ In the medical field, disrupting biofilm cohesion is a standard procedure in chronic wound care, making the remaining adherent bacteria more susceptible to antibiotics.^2^ Understanding biofilm mechanics is also essential for preventing biofilm formation or for leveraging their benefits in industrial and bioprocessing applications, such as wastewater treatment.^3^ Broadly, research on biofilm mechanics aims to deepen our understanding of the biofilm life cycle,^4^ clarify how the viscoelastic properties of biofilms aid bacteria survival in mechanically challenging environments (e.g., under flow)^5^ and inspire new approaches for the mechanical removal of biofilms.^6^

Biofilms are often described as complex fluids with viscoelastic material properties^7^ or as composite materials, where relatively rigid colloid-like bacteria are embedded within a soft, hydrated ECM composed of various biopolymers.^8^ The ECM accounts for a significant volume fraction of the dry biomass so that the nature of the biopolymers, their molecular structure as well as their interactions are expected to govern the biological material responses at macroscopic scales.^9,10^ The elastic modulus (also called Young’s modulus or stiffness) is a mechanical parameter broadly used to report on the rigidity of a material, e.g., the elastic modulus of bone is about 20 GPa and 1000 times higher compared to skin (20 MPa). In cystic fibrosis, the *Pseudomonas aeruginosa* bacteria that colonize lungs increase biofilm elastic modulus by increasing their production of the ECM polysaccharide Psl, whereas an increased expression of Pel and alginate has no influence on the biofilm stiffness.^11^ Similarly, *Pseudomonas fluorescens* biofilms become more ductile (less brittle) when the ECM-to-cell ratio is increased.^12^ Recently, it was found that the molecular structure of curli amyloid fibers extracted from *E. coli* biofilms depends on the growth conditions and correlates with biofilm stiffness.^13^ Interactions between the ECM components produced by *Streptococcus mutans* were also shown to promote biofilm cohesion and prevent mechanical biofilm failure when scraping with the tip of an indenter.^14^

Biofilm structural heterogeneity contributes to its mechanical complexity. Gradients of nutrients and oxygen, emerging across the biofilm, result in gradients of metabolic activities and ECM production.^7^ In the macrocolony model, for example, cross-sections of *E. coli* biofilms, grown on the surface of nutritive agar plates, revealed layers with distinct ECM content and organization.^15^ The mechanical stability of such *E. coli* AR3110 macrocolony biofilms was proposed to benefit from the assembly of a curli and phosphoethanolamine-cellulose (pEtN-cellulose) fiber network, which provides tissue-like elasticity.^16,17^ The small number of studies reporting on the mechanical properties of *E. coli* biofilms did, however, not specifically assess the mechanical implications of such an elaborate architecture.^18^ Macrocolony biofilms from various species were also shown to exhibit porous structures or channels for nutrient transport and diffusion.^19–21^ It is expected that local composition and architecture determine biofilm mechanics at the microscopic scale. As a result, the mechanical properties are highly heterogenous throughout the biofilm and differ significantly from the macroscopic properties.^22^ Due to the living nature of biofilms, their mechanical properties further depend on the time scale considered (e.g., short time scales accounting for ECM rearrangement or long time scales related to growth phenomena or changes in gene expression).

To characterize the viscoelastic properties of macrocolony biofilms, the choice of method as well as the accessible time and length scales are essential aspects to consider.^1^ Oscillatory shear-rheology has been widely used to assess the bulk viscoelastic properties of bacterial macrocolony biofilms produced by *Pseudomonas spp*.,^23^ *Azotobacter vinelandii,*^24^ and *Vibrio cholera.*^25^ While this method gives information at the macroscopic scale, the sample preparation often involves scraping the biofilms from their substrate, potentially destroying their ECM architecture.^26^ An alternative is to grow biofilms directly on the rheometer sample stage, as demonstrated with several strains of *Bacillus subtilis* and *Staphylococcus epidermidis*.^27^ For measuring local properties of native macrocolony biofilms, atomic force microscopy (AFM) is attractive as it provides height and stiffness maps with single-cell spatial resolution.^28^ Microindentation may be better suited for microscale mechanical characterization (as opposed to nanoscale). On the macrocolony scale (centimeter-sized), indenters with a diameter of tens of micrometers provide local information while still averaging over the contribution of several tens of bacteria and the surrounding fibrous ECM network. For example, microindentation showed that growing *E. coli* on drier agar substrates yields stiffer biofilms.^29^ Modelling stress relaxation of biofilms in rheology experiments revealed multiple characteristic time scales as a function of ECM composition.^30^ Local biofilm viscoelastic properties can also be obtained from passive or active micro-rheology, i.e., by tracking the Brownian motion or induced displacement (with magnetic or optical tweezers) of micro-particles embedded in the ECM.^26^ This last approach is preferred for immersed biofilms grown in liquid, e.g., in flow chambers.^31,32^ Overall, the diversity of mechanical tests available, the lack of standardization in the protocols for biofilm characterization and the complex response of biofilms to deformations lead to difficulties in comparing mechanical parameters across different studies,^26^ even when considering the same bacterial strain grown in different^1^ or very similar conditions.^33^

In this work, we investigated the contributions of the main ECM components of *E. coli* biofilms, namely curli amyloid and pEtN-cellulose fibers, to the mechanical properties of the macrocolony biofilm material. Moreover, we explored the relevance of their interactions as well as the integrity of their architecture for the viscoelastic behavior of the biofilm. We grew different *E. coli* K-12 mutants on nutritive agar substrates to obtain macrocolony biofilms of different ECM compositions, and characterized their viscoelastic properties using oscillatory shear-rheology and microindentation. For rheology, the biofilm material was scraped from the agar substrate and homogenized while the native biofilm architecture was preserved when performing microindentation. Comparing the two characterization methods reflects the difficulties mentioned above.^26^ At the same time, the results indicate that curli amyloid fibers play a key role in determining biofilm rigidity while cellulose, especially pEtN-cellulose, contributes to maintaining its structural integrity. Further structural investigations using confocal microscopy suggest that the composite nature and structural heterogeneity of the macrocolony biofilms make them sensitive to the specific methods used for mechanical characterization. Notably, this sensitivity is influenced by the composition of the ECM. All in all, this work does not only bridge an important gap in the mechanical characterization of *E. coli* macrocolony biofilms. It also reminds the biofilm community that multi-modal mechanical characterization is needed to understand the mechanical properties of biofilm-based materials.

## Results

### Influence of EPS synthesis on macrocolony biofilm morphology, texture and mass

To investigate the influence of the *E. coli* ECM composition on biofilm growth and material properties, we cultured wild-type and mutant strains with varying capacities to produce ECM components for 7 days on salt-free LB agar substrates. The different strains produce neither curli nor cellulose (AR198, also referred to as “No ECM”), only non-modified cellulose (AP472), only pEtN-cellulose (AP329), only curli (W3110), curli and non-modified cellulose (AP470), and finally both curli and pEtN-cellulose fibers (AR3110). Moreover, we grew biofilms from a bacterial suspension containing a 1:1 mixture of W3110 and AP329 so that both curli and pEtN-cellulose should be present but produced by different bacteria (Fig. 1A).

**Figure 1.**
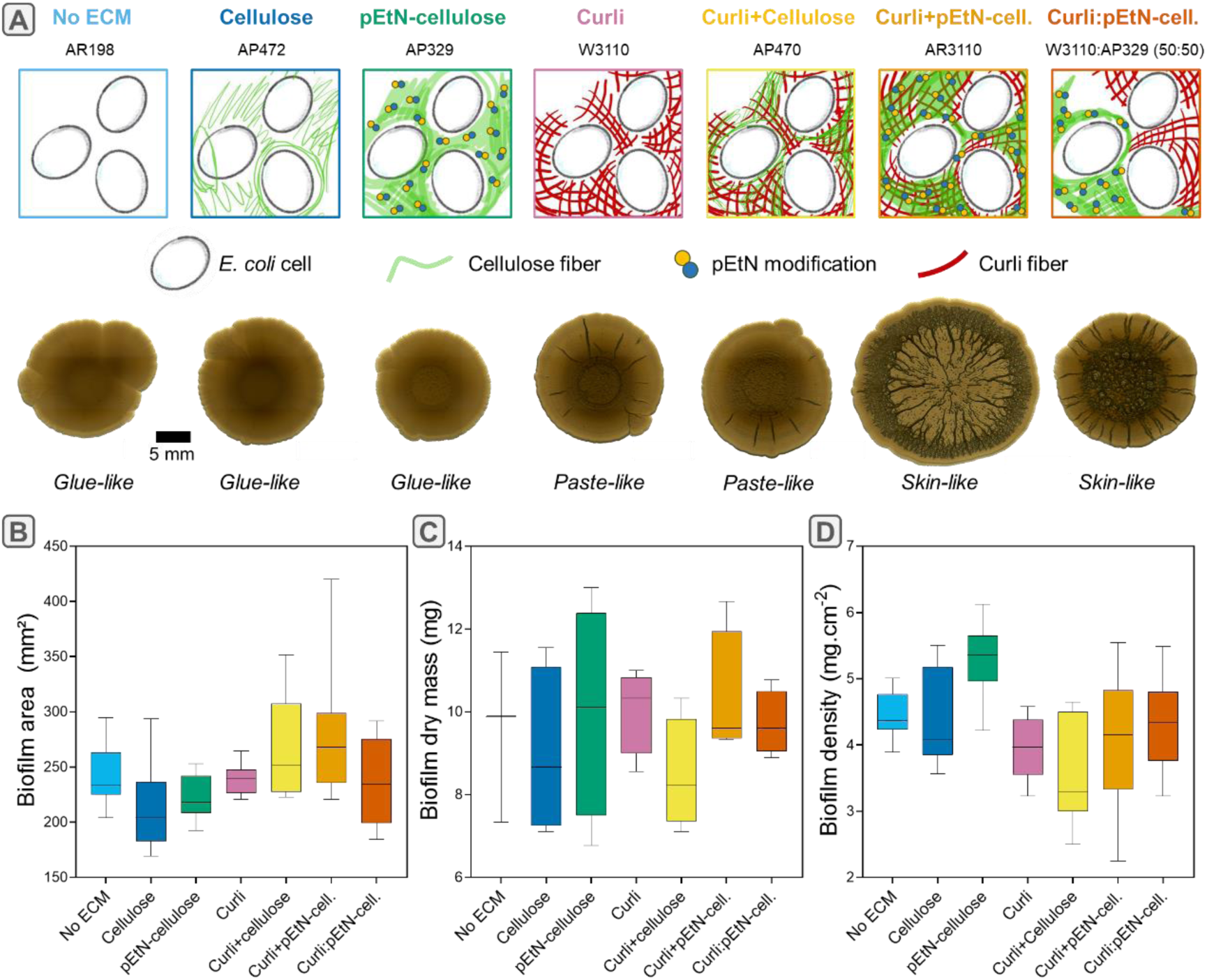
ECM composition influences macrocolony biofilm spreading, texture, dry mass and water content. (A) Morphologies and textures of biofilms grown from *E. coli* strains with various ECM compositions. The images were acquired after 7 days of growth using a stereomicroscope in transmission. The darker lines result from the emergence of wrinkles in the third dimension. (B-D) Biofilm diameter (n=12 biofilms produced in 4 different plates), dry mass (N=4 plates per strain) and density as a function of composition; median, quartiles and extreme values are represented by the line, the box limits and the whiskers respectively; Mann-Whitney U-tests were performed for statistical analyses. Statistical significance values are reported in Tables S1-3.

As reported before, ECM composition greatly influences macrocolony biofilm morphology and texture.^34,35^ We observed that biofilms grown from curli-producing bacteria present apparent wrinkles, while others do not. Biofilms containing both curli and pEtN-cellulose form numerous and predominantly radial delaminated wrinkles independent on whether these fibers are produced by the same (AR3110) or different bacteria (W3110:AP329). Under our experimental conditions, only these two cases yielded biofilms with a skin-like texture upon handling, which demonstrates some cohesion in the material. In contrast, biofilms containing only curli fibers (W3110) or curli fibers and non-modified cellulose (AP470) formed fewer and shallower radial wrinkles. They exhibited paste-like properties, quickly losing the film aspect upon manipulation, while retaining some cohesion (Fig. 1A). Biofilms lacking the curli component, i.e., producing only pEtN-cellulose (AP329), only non-modified cellulose (AP472) or none of the ECM fibers (AR198), all appear featureless and exhibit glue-like textures. These biofilms adhered strongly to the spatula when scraped from the soft agar substrate.

Biofilms containing both curli and pEtN-cellulose (AR3110) were slightly more spread out than their counterparts after 7 days of growth (Fig. 1B and Table S1) while biofilms containing only non-modified cellulose spread slightly less. However, all seem to contain similar amounts of dry mass, i.e., between 7 and 11 mg, with slight but statistically non-significant variations from strain to strain (Fig. 1C and Table S2). This yields different biofilm surface densities (in mg cm^-1^), accounting for how much biofilm dry mass has formed on a specific area of the substrate (Fig.1D). Even though no apparent macroscopic wrinkles were present, biofilms containing only pEtN-cellulose showed a significantly higher surface density than the other strains (Table S3). Water content and water activity, measured in 7-day old biofilms, did not show statistical differences between strains (Fig. S1). It is yet important to highlight that the low masses measured introduce measurement errors that cannot be easily offset by increasing the number of technical repeats across all conditions.

### Influence of ECM composition on the bulk viscoelastic properties of homogenized biofilms measured by oscillatory shear-rheology

To probe the viscoelastic properties of biofilms with different ECM composition, we performed bulk oscillatory shear-rheology using so-called “homogenized biofilms”. For each *E. coli* strain, a few biofilms grown in the same Petri dish were scraped off the surface and mixed together to collect sufficient material to fill the 250 μm gap between the two parallel plates in the rheometer setup (Fig. 2A). Amplitude sweeps were performed to determine the viscoelastic response of the homogenized biofilms to the applied shear strain (Fig. 2B). For each ECM composition, the plateau values of the storage modulus (G’_0_) were almost one order of magnitude larger than those of the loss modulus (G"_0_). Comparing the median values of G’_0_ and G"_0_ for the different *E. coli* strains revealed different viscoelastic responses (Fig. 2C and D). For example, homogenized biofilms containing only pEtN-cellulose or curli fibers presented storage moduli of 18 and 16 kPa, respectively. In contrast, biofilms containing both ECM components were stiffer, namely 28 kPa when pEtN-cellulose and curli were produced by the same bacteria (AR3110; Curli+pEtN-cellulose) and 51 kPa when they were synthesized from the mixture of W3110 and AP329 that each produces one type of fiber (Curli:pEtN-cellulose). Interestingly, homogenized biofilms containing only non-modified cellulose also presented a storage modulus of 51kPa and biofilms producing neither fiber were not the softest (G’_0_ = 31 kPa).

**Figure 2.**
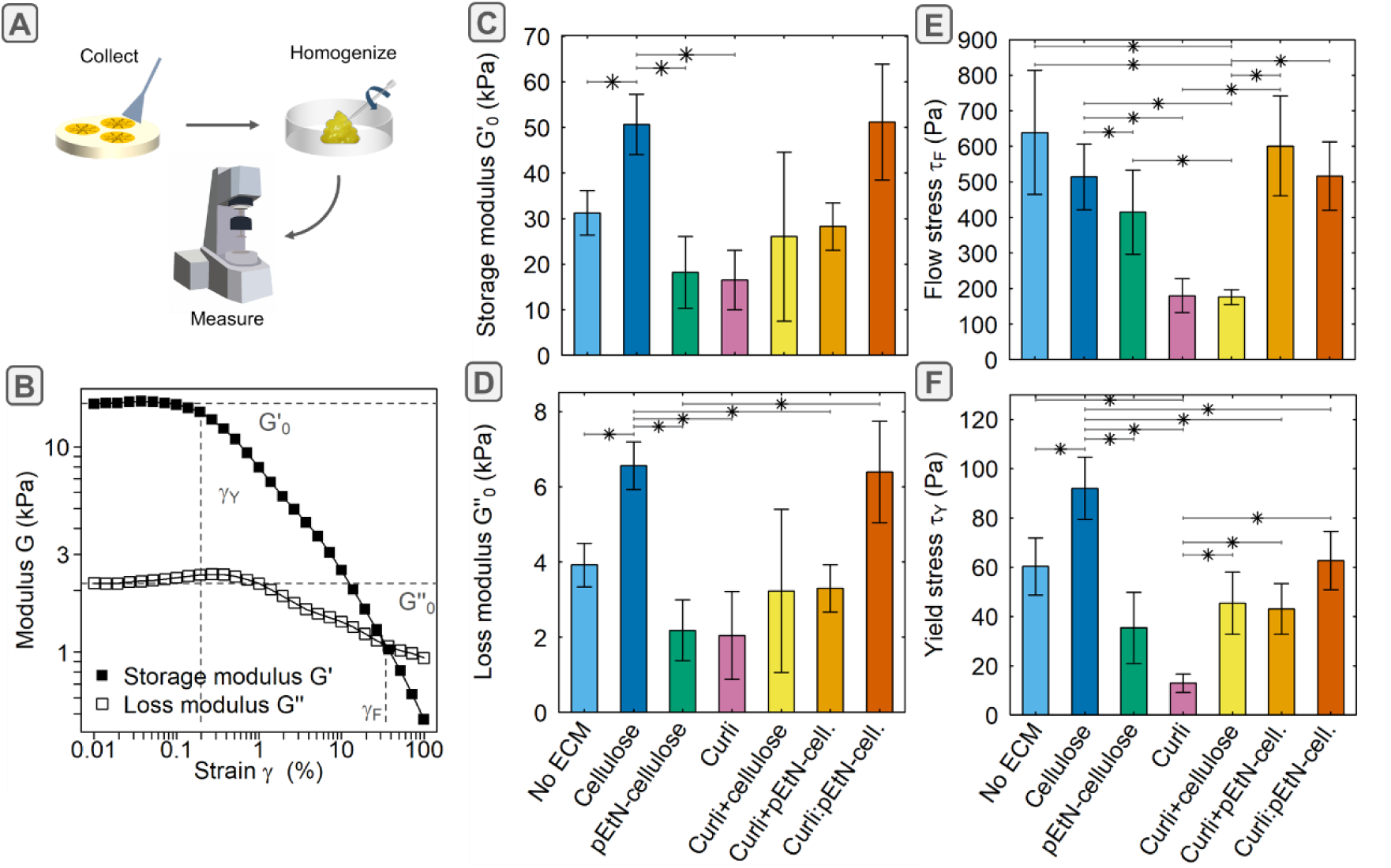
Oscillatory shear-rheology of *E. coli* biofilms with different ECM compositions. (A) Schematic image of sample preparation steps for bulk rheology measurements of homogenized biofilms. (B) Representative strain amplitude sweep (here for AR3110) used to extract the viscoelastic properties of homogenized biofilms (ω = 10 rad s^-1^, γ = 0.01 to 100%). (C) Plateau of the storage modulus (G’_0_). (D) Plateau of the loss modulus (G"_0_). (E) Yield stress (τ_Y_). (F) Flow stre_s_s (τ_F_). (C-F) Bars represent median values and error bars represent standard deviation of bootstrapped data sets. Α Wilcoxon signed rank test was used (*: p-value=0.1, n=3). More details are given in Materials and Methods.

The yield stress (τ_Y_) was also determined from the obtained amplitude sweeps as the product of the yield strain (γ_Y_) and the shear modulus 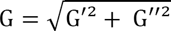 (Fig. S2). The yield stress represents the limit of the linear viscoelastic range, i.e., the amount of stress where irreversible plastic deformation begins to occur. Biofilms producing only curli displayed a lower yield stress than their counterparts where curli are associated with (pEtN-)cellulose (Fig. 2E). Interestingly, the biofilms containing both fibers yielded at a stress below 43 Pa for AR3110 and 62 Pa for the mixture of W3110 and AP329. Biofilms lacking both fibers showed a yield stress of 60 Pa and biofilms producing only non-modified cellulose resisted yielding up to 92 Pa, which is higher than any other ECM composition.

The flow stress (τ_F_) was derived as the flow strain (γ_F_) multiplied by the shear modulus 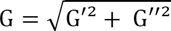 and corresponds to the stress where the viscous component begins to dominate over the elastic component (G’ < G"). At this point, more energy is irreversibly dissipated than reversibly stored and the material predominantly behaves like a fluid. Biofilms containing only curli or curli in association with non-modified cellulose started to flow at 180 Pa. This was lower than homogenized biofilms with any other ECM composition that began to flow at stresses larger than 400 Pa (Fig. 2F).

### Influence of ECM composition on the local mechanical properties of native macrocolony biofilms measured by microindentation

In contrast to oscillatory shear-rheology, which requires harvesting and homogenizing several biofilms for one bulk measurement, microindentation enables several local experiments on a single biofilm while preserving its native architecture. To assess the contribution of different ECM components and their architecture to the viscoelastic properties of the biofilms, we complemented our mechanical characterization with microindentation experiments in the center of intact biofilms. For each *E. coli* strain, biofilms for microindentation were grown in similar conditions as for the rheology experiments. Microindentation was performed with a 50 µm diamond indenter in the “air-indent” mode.^36^ In this mode, the measurement starts before reaching the biofilm so that the detection of the surface upon contact is facilitated after experiencing a slightly negative force (Fig. 3A) (displacement δ = 0). Further tip penetration constitutes the loading phase. At the maximum indentation, a holding step of 10 s was set to measure the viscoelastic relaxation of the compressive stresses. Finally, the tip was retracted to the initial position far above the surface, while the corresponding adhesive forces were recorded during the detachment phase.

**Figure 3.**
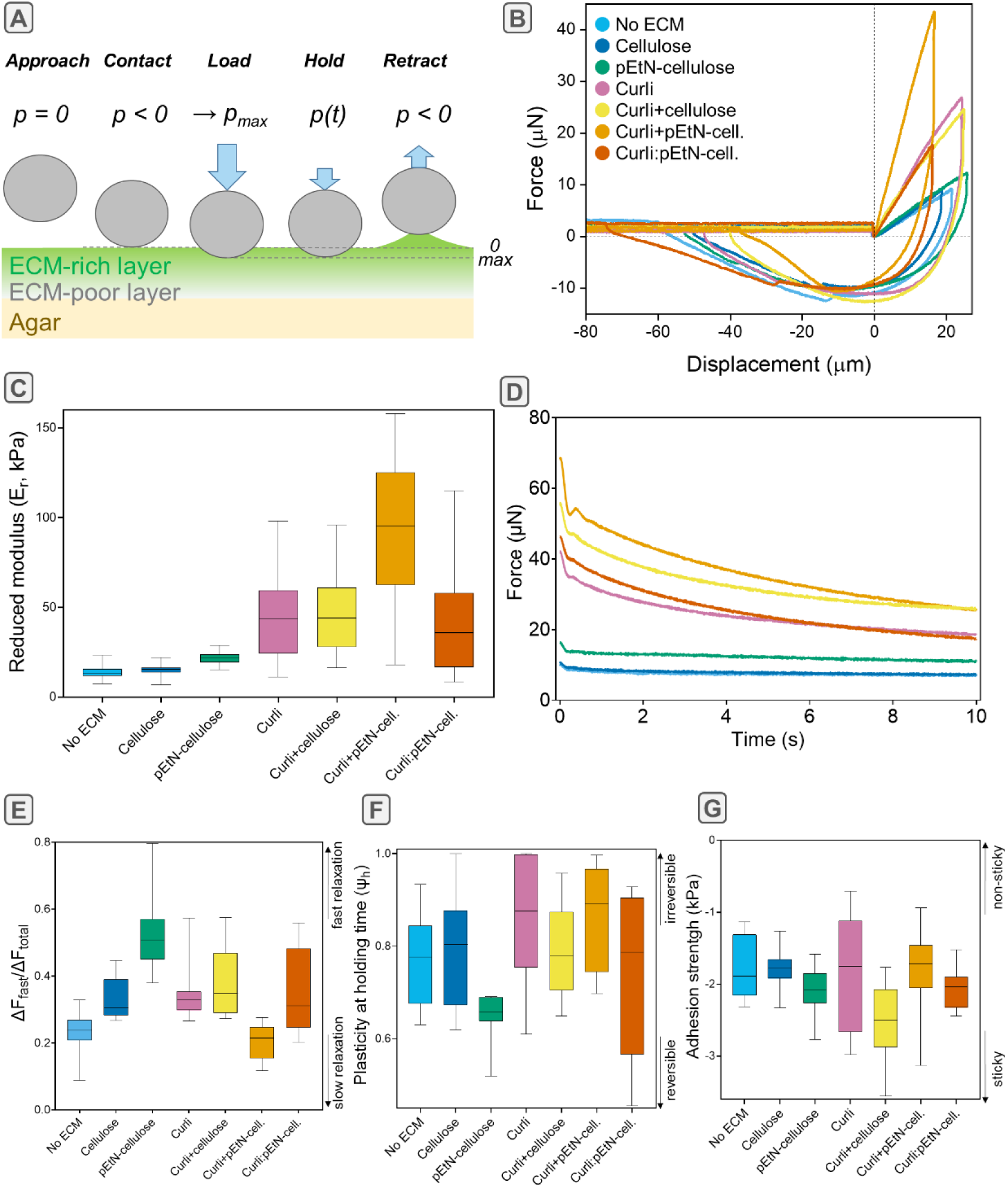
Mechanical characterization of *E. coli* macrocolony biofilms with microindentation. (A) Schematic image of the microindentation experiment performed on the native biofilms. (B) Representative force-displacement curves of *E. coli* biofilms with different ECM compositions, after the contact point is set at (0;0) for further analysis. (C) Reduced elastic moduli *E*_r_. (D) Representative force-time curves of each strain during the holding time at a maximum penetration depth of 20 µm. (E) Portion of the force relaxation happening before the tip instability *ΔF*_fast_/*ΔF*_total_, derived from the relaxation curves. (F) Plasticity during the holding time, defined as the ratio of energy dissipated during relaxation and energy stored upon indentation. (G) Adhesion strength σ_Adh_ measured from the minimum force recorded during tip retraction divided by the contact area at maximum indentation. Data in E-G are derived from indentation curves with a maximum depth penetration of 20 µm. See methods section and Figure S3 for technical details. Median, quartiles and extreme values are represented by the thick line, the box limits and the whiskers respectively; n = 10 curves from N = 4 different biofilms. Statistical analysis was performed with a Mann-Whitney U test and is reported in Tables S4-7.

The slopes of the load-displacement curves characterize the stiffness of the biofilm material upon compressive loading (See Fig. S3A and Methods). Figure 3B shows that all biofilms producing curli reached higher maximum loads (p_max_) than curli-deficient strains when loaded to comparable indentation depths. The reduced elastic moduli (*E*_r_) were determined, fitting a Hertzian model for small displacements in the linear elastic region (<10 µm) (Fig. 3C). The stiffest biofilms were those containing pEtN-cellulose and curli produced by the same bacteria (AR3110; 97 kPa), followed by the other biofilms containing curli (W3110, AP470, 1:1 mixture of W3110 and AP329) with values around 40 kPa. These values are about twice as high as those measured for all curli-deficient biofilms (Fig. 3C). Statistical analyses suggested no difference between biofilms containing non-modified cellulose or no ECM (Table S4). No significant differences were further observed between the mixed biofilm and biofilms obtained from the individual strains (W3110 or AP329) as well as between biofilms containing both non-modified cellulose and curli produced by the same bacteria (AP470). These results indicate that the pEtN modification critically determines biofilm compressive stiffness.

As for many soft living materials, the mechanical properties of *E. coli* biofilms are expected to be time-dependent. To determine how their ECM composition and architecture affect their viscoelastic properties, relaxation was measured while holding the indenter for 10 s at the maximum displacement (Fig. 3D). A local load maximum, consistently observed at 350 ms (Fig. S3C), was attributed to a tip instability between 250 and 350 ms as the instrument switched from displacement-controlled indentation to constant displacement for the holding phase. Fast relaxation may be attributed to the flow of water in the porous network (< 1 s) and slow relaxation to the reorganization of the ECM network itself (< 10 s).^18^ We thus assessed the fast relaxation behavior by measuring the portion of force relaxed until the tip instability with respect to the total amount of force relaxed within the 10 s holding time (*ΔF*_fast_/*ΔF*_total_) (Fig. 3D,E and Table S6). Most of the biofilms relaxed about 30 to 35% of the force before the tip instability. However, biofilms containing only pEtN-cellulose dissipated more than 50% of the total relaxed force in this short time window. In contrast, biofilms containing both curli and pEtN-cellulose formed by the same bacteria (AR3110) appeared to relax only 20% of the force in the short time window, thus suggesting that the relaxation is rather dominated by ECM rearrangement. Bacterial aggregates without ECM also showed a lower portion of force relaxation within the first 205 ms; here, most probably due to bacteria rearrangement. Note that such characteristic times remain short compared to biofilm growth and bacterial reorganization dynamics (> 100 s),^37^ which are thus expected to play a minor role.

To compare the energy dissipated in different biofilms, we defined the plasticity during the holding time (Ψ_h_) as the ratio of the energy dissipated during relaxation and the total energy stored upon indentation (see Materials and Methods). While all biofilms dissipated more than half of the energy stored (Ψ_h_>0.5), biofilms containing only pEtN-cellulose (AP329) consistently dissipated less compared to biofilms with other ECM compositions (Fig. 3F and Table S6). Biofilms formed by the mixed strains were an exception but presented a broad distribution of plasticity values. In contrast, biofilms containing only curli (W3110) dissipated almost all energy during the holding time (Ψ_h_ = 0.93), which was similar to the energy dissipated by biofilms producing both curli and pEtN-cellulose formed by the same bacteria (AR3110) (Ψ_h_ = 0.85) but higher than for the other biofilms (see Table S6 for statistical significance values).

During the retraction phase, negative forces (p < -10 µN) and displacements (< -70 µm) were necessary to detach the tip from the biofilm surface. Values characterizing the adhesion strength were derived from the minimum forces measured during retraction divided by the maximum contact area between the indenter and the biofilm (i.e., at maximum displacement). While the broad distributions of adhesion strengths do not allow for clear conclusions, statistical analysis suggests that biofilms containing both curli and non-modified cellulose were more adhesive than the others with an adhesion strength of 2.50 kPa (Fig. 3G and Table S7). Moreover, biofilms with ECM made of pEtN-cellulose only appeared to adhere more than biofilms containing non-modified cellulose only (2.08 kPa vs 1.79 kPa). This is another indication that pEtN-cellulose has an important contribution to the mechanical biofilm properties.

### Importance of intact macrocolony biofilm morphology, ECM composition and fiber arrangement

Microindentation showed a broad distribution of viscoelastic properties and the trends observed with this method did not always reflect those measured with bulk shear-rheology. To better understand these discrepancies, we first investigated the major structural differences between the various *E. coli* biofilms. We then focused on biofilms from *E. coli* producing curli and/or pEtN-cellulose and aggregates from *E. coli* producing no ECM to study how the respective biofilm architectures were affected by a homogenization step (mixing).

As previously shown, biofilm internal architecture highly depends on ECM composition.^35^ Here, fresh biofilms grown on salt-free LB agar, supplemented with Direct Red 23 (also known as Pontamine Fast Scarlet 4B), were sandwiched using a second layer of agar. Sections with a thickness of ∼ 1 mm were cut in the vertical direction along the biofilm diameter (+/- 1 mm precision), and the resulting cross-sections were imaged with confocal microscopy (Fig. 4). The obtained images provide a better understanding of differences in biofilm wrinkling and also reveal the internal organization of the ECM. As expected, samples from bacteria producing no ECM barely display a fluorescent signal. In agreement with the stereomicroscopy images (Fig. 1A), biofilms containing only non-modified cellulose were rather flat and biofilms containing both curli and pEtN-cellulose displayed many wrinkles with high aspect ratio (Fig. 4). However, while biofilms containing only pEtN-cellulose appeared flat from above, the cross-sections revealed a lot of small, thick and densely packed wrinkles, consistent with what has been previously described.^16^ This observation also agrees with their higher surface density (Fig. S4). For biofilms containing only curli, very few wrinkles were seen both with stereomicroscopy and in the cross-sections. Cross-sections of biofilms containing curli and non-modified cellulose displayed more wrinkles than biofilms containing only curli, but less than biofilms containing both curli and pEtN-cellulose. Further investigation of the microstructure of the stained ECM showed that the ECM appears granular in biofilms containing non-modified cellulose only, curli only and the combination of both, while biofilms containing pEtN-modified cellulose showed a fibrous ECM with layers.

**Figure 4.**
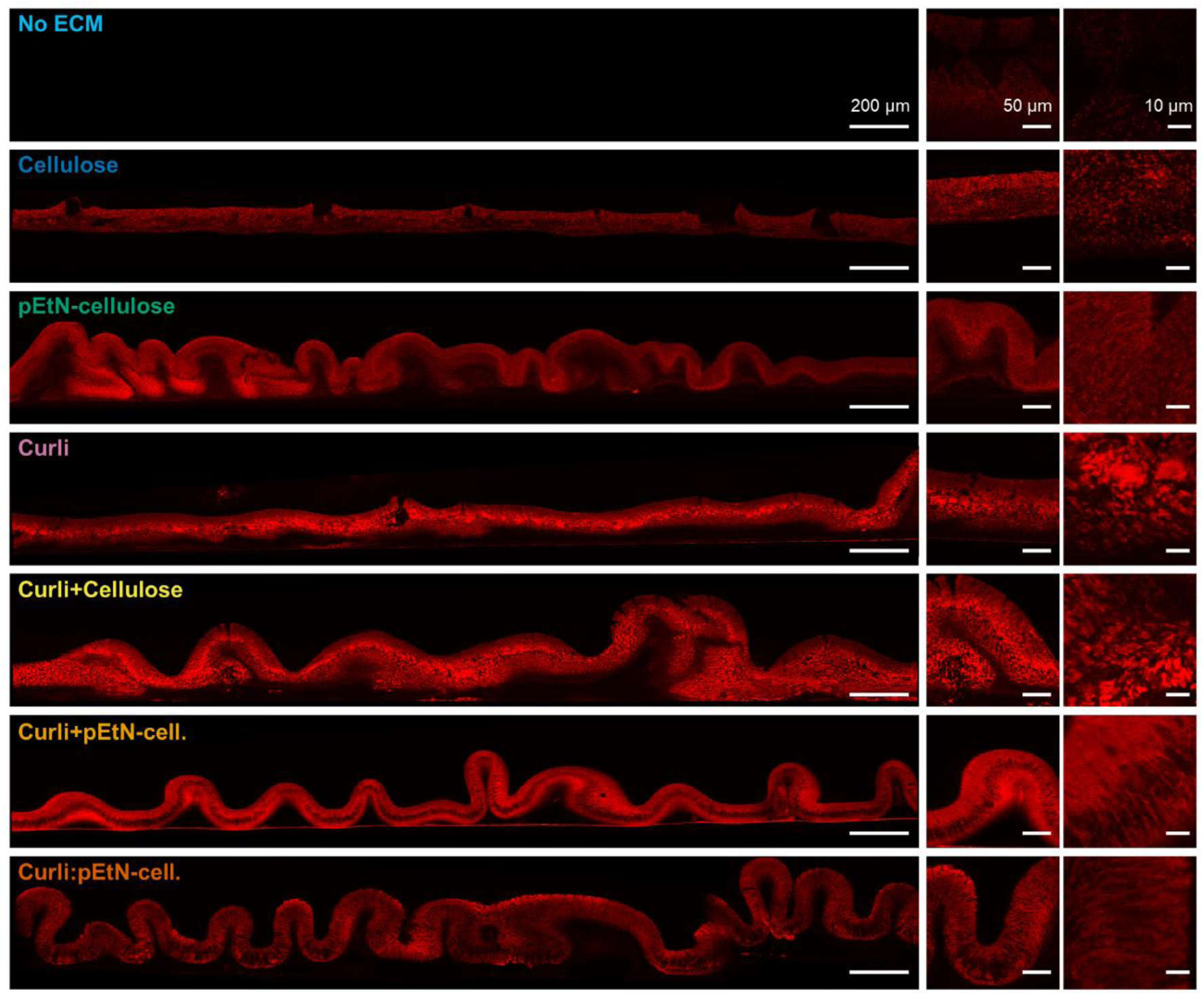
Structure of macrocolony biofilms grown from the different *E. coli* strains. Confocal images of cross-sections of native biofilms grown at 28 °C for 7 days on salt-free LB agar supplemented with Direct Red 23 to stain the ECM (red). The images were acquired with 552 nm laser excitation and photon collection in the range from 600 to 700 nm. All images are displayed with the same brightness and contrast settings.

One major difference between oscillatory shear-rheology and microindentation lies in the homogenization step required for bulk rheology. To assess the structural effect of biofilm scraping and mixing, the same imaging procedure was performed on homogenized biofilms, prepared as for the rheology experiment. Figure 5A presents a direct comparison of images taken for native *vs.* homogenized biofilms, now focusing on bacteria that produce no ECM, only pEtN-cellulose, only curli or both of these ECM fibers. No structural change was visible when mixing biofilms containing none of the fibers. The densely packed patches observed in native biofilms containing curli only appeared to be more evenly distributed in the homogenized mass. The wrinkly morphology of biofilms containing pEtN-cellulose or pEtN-cellulose in combination with curli fibers was greatly affected by the mixing step. After mixing, the overall structure was destroyed and the folds were loosely pressed against each other. At the same time, the layer structure within the biofilms remained visible, suggesting partial homogenization of the macroscopic structure while the microscopic structure remained intact.

**Figure 5.**
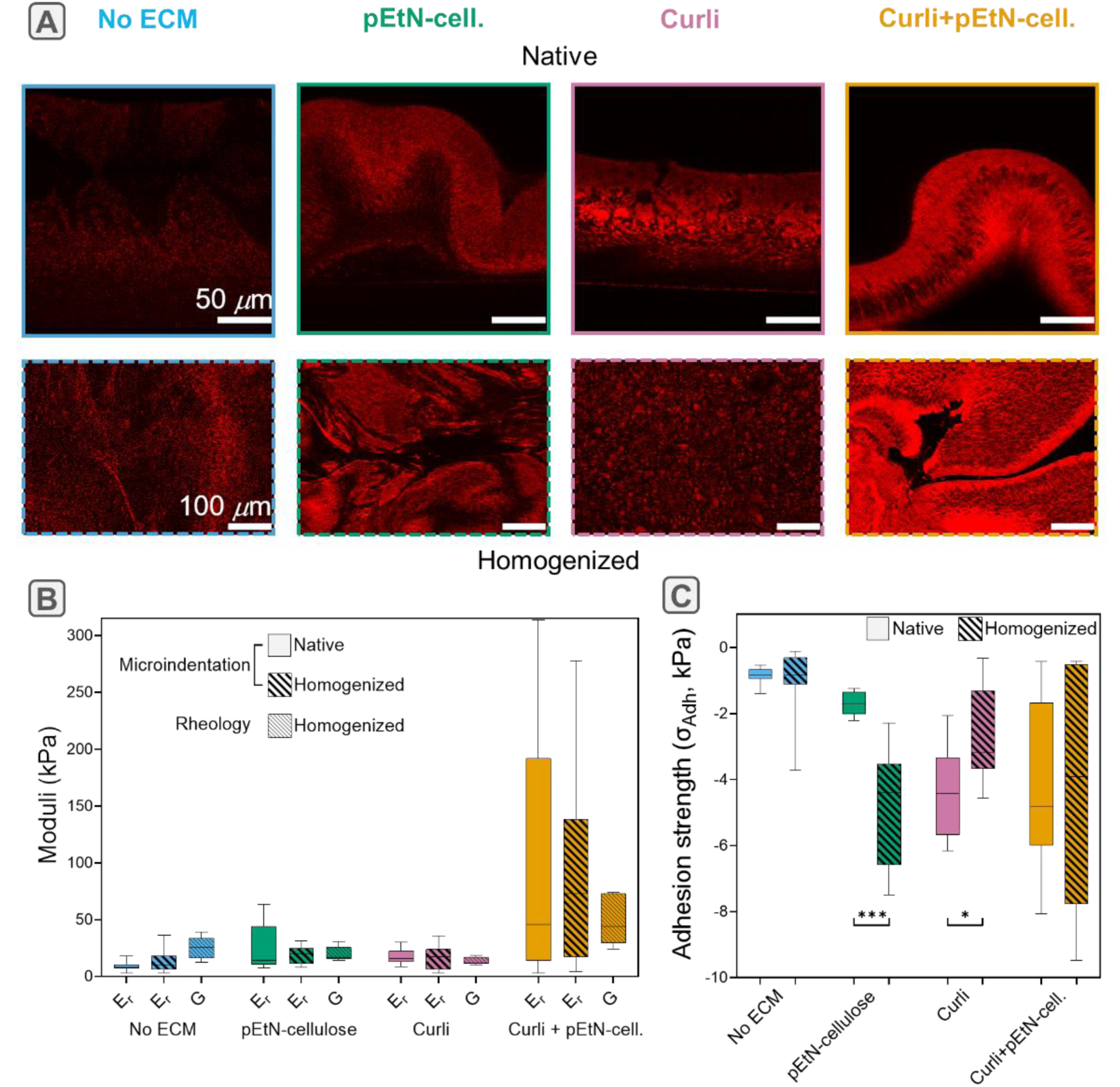
Influence of a homogenization step (mixing) on the structure and mechanics of selected *E. coli* macrocolony biofilms with different ECM composition. (A) Confocal images of native *vs.* homogenized biofilms, formed by bacteria producing no ECM, only pEtN-cellulose, only curli or both ECM fibers. (B) Comparison of the reduced indentation elastic moduli E_r_ of native and homogenized biofilms with the shear modulus G obtained from rheology. (C) Adhesion strength σ_Adh_ measured from the minimum load recorded during retraction. The values are normalized by the contact area at maximum indentation. Median, quartiles and extreme values are represented by the horizontal line, the box limits and the whiskers respectively; n = 7 curves from N = 3 different biofilms. Statistical analysis was performed using a Mann-Whitney U test and are significance values reported in Table S8.

To assess how structural changes upon homogenization affect biofilm mechanical characterization, we directly compared oscillatory shear-rheology and microindentation with biofilms grown on the same plates. This experimental design minimizes the contribution of plate-to-plate variability, frequently observed when growing biofilms. The compressive and shear moduli measured in this comparative experiment followed similar trends as those obtained with the two independent measurement campaigns described above. The median of the reduced indentation elastic moduli of native biofilms (i.e., conserving their original architecture) were the lowest for curli-deficient biofilms (∼ 7.9 to 13 kPa), slightly higher for curli-only biofilms (18 kPa) and significantly higher for biofilms containing both curli and pEtN-cellulose (26 kPa) (Fig. 5B and Table S9). The rheology experiments showed a different trend. Homogenized samples containing no ECM or biofilm with both curli and pEtN-cellulose presented higher shear moduli (25 and 44 kPa, respectively) than biofilms with only pEtN-cellulose or curli (16.4 and 11.6 kPa, respectively).

To explore the role of sample preparation, this experiment also included microindentation on homogenized biofilms. Different mechanical parameters were compared between native and homogenized biofilms containing the different ECM compositions. No statistical difference was observed between the reduced moduli (Fig. 5B and Table S9), the plasticity at holding time (Fig. S5A and Table S9) nor the portion of force relaxation happening fast (Fig. S5B and Table S9). However, a twofold increase in adhesion strength was observed for homogenized biofilms containing only pEtN-cellulose, as well as a slight decrease for biofilms containing only curli (Fig. 5C and Table S9).

Figure 6 represents the shear moduli obtained from oscillatory shear-rheology (i.e., after sample homogenization) plotted against the reduced elastic moduli obtained with microindentation. Figure 6A shows data compiled from the shear-rheology and microindentation experiments reported in Figures 2 and 3. These experiments were performed independently, i.e. not at the same time and by different operators. Figure 6B shows data from comparative experiments where biofilms were grown on the same plate. Three plates were analyzed per selected *E. coli* strain. The dashed lines represent the theoretical relation between the shear modulus G and the reduced modulus 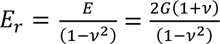 for ideal materials with different Poisson’s ratios.

**Figure 6.**
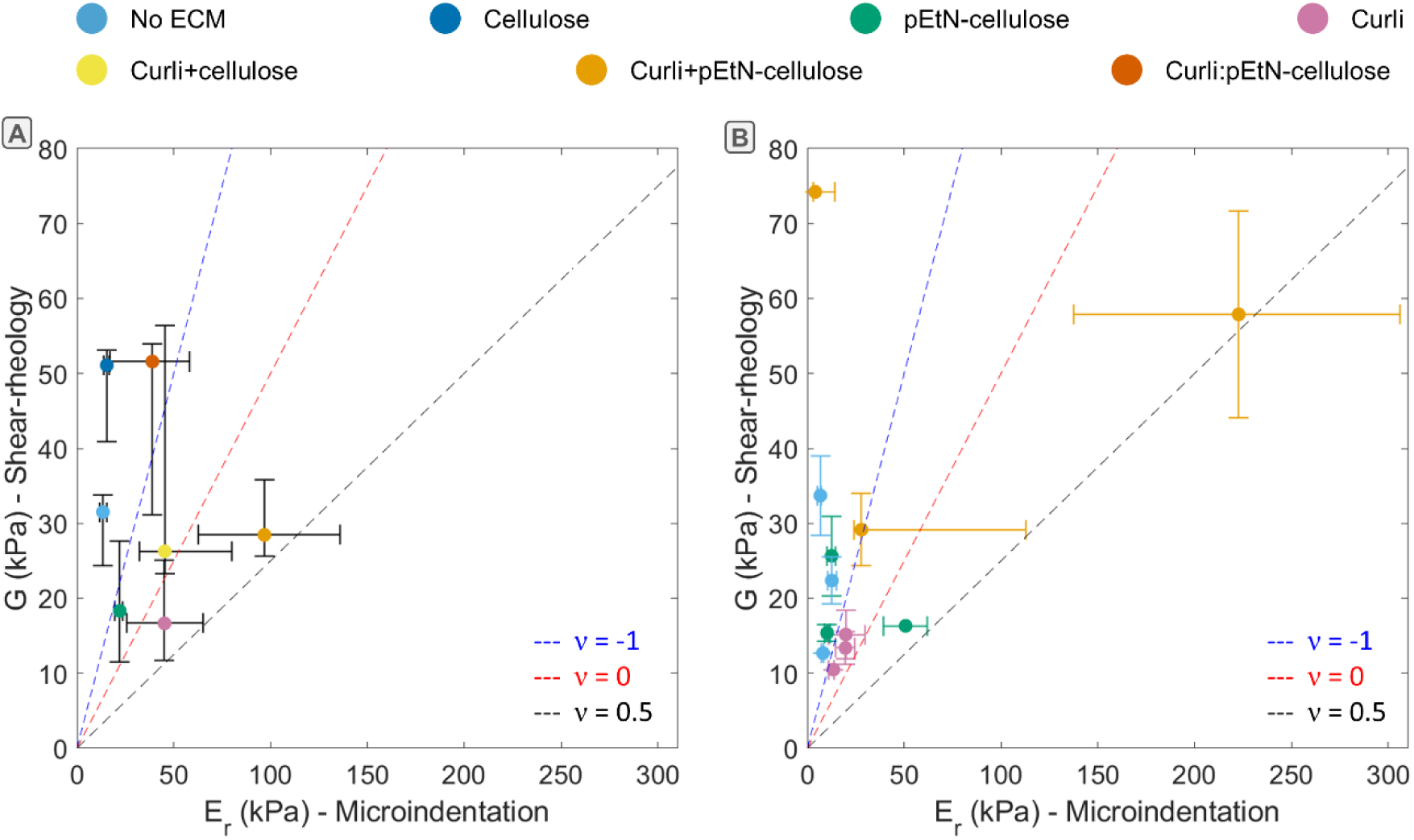
Shear modulus *vs* reduced elastic modulus obtained on macrocolony biofilms with different ECM compositions. Each point represents median values. The error bars show the 0.25 and 0.75 quantiles of the data distribution. (A) The data are compiled from the independent experiments reported in Figures 2 and 3. (B) The data correspond the comparative experiments performed with biofilms grown on the same plate (N=3 plates, n=3 biofilms per plate). The dashed lines represent theoretical relations between the shear modulus G and the reduced modulus E_r_ for ideal materials with different Poisson’s ratios.

The scattered positions of the experimental points with respect to these lines confirm that biofilms are complex materials with non-trivial mechanical behavior. This result also suggests that the experiments become hardly comparable to the theory after modifying biofilm internal structure through homogenization. Another observation is the wide distribution of moduli, both for the rheology and the microindentation data, despite limiting plate-to-plate variations. This observation was especially striking for data obtained for biofilms grown from bacteria producing both curli and pEtN-cellulose. In contrast, biofilms deficient in curli seem to yield narrower distributions of reduced moduli obtained by microindentation. Aware of possible variations in agar properties from plate to plate, we verified that there was no systematic effect of the agar reduced modulus on biofilm reduced modulus (Fig. S6). Despite the large distribution of values, biofilms containing none or only one type of fiber tend to have low moduli (the corresponding points are located at the bottom left of Fig. 6), whereas biofilms containing curli and (modified-)cellulose tend to have larger moduli (top right).

## Discussion

This work reports on the mechanical analysis of macrocolony biofilms formed by a collection of *E. coli* strains that produce ECM of different compositions. Comparing oscillatory shear-rheology and microindentation, we found that microindentation yields more conclusive results. Indeed, microindentation, which conserves the native structure of the biofilms, indicates that biofilms containing curli fibers tend to be stiffer and that the pEtN-modification of cellulose enhances biofilm stiffness further when curli fibers are present.

Oscillatory shear-rheology and microindentation are two distinct mechanical characterization methods that fundamentally differ in two key aspects. Rheology applies a shear stress to the sample, whereas microindentation predominantly probes it under compressive stress. Additionally, bulk rheology averages the properties on one entire or several biofilms, while microindentation locally probes the biofilm over a region that encompasses numerous bacteria and their surrounding ECM (∼3000 to 7000 µm²). A quantitative comparison of the shear and compressive moduli requires knowledge of the Poisson’s ratio, which is not easily available for such highly heterogeneous materials. However, biofilms are not necessarily expected to exhibit similar mechanical responses under these different conditions and at the scales analyzed. This highlights the complementarity of the two methods, especially for investigating the differential mechanical responses of biofilms grown from different strains, as demonstrated in this work.

The sample preparation strategies chosen are also very different for the two methods. It is in principle possible to transfer intact macrocolony biofilms from the agar plate directly to the rheometer; however, the heterogeneity of the sample is rarely taken into account and biofilm wrinkles are compressed while reaching the final gap size.^38^ In the literature, homogenization steps are scarcely applied to reduce potential variability due to a random positioning of the biofilm structure when scrapping and transferring the sample from the agar to the rheometer.^39^ Here, we gently mixed the samples to obtain a homogenized and compact biofilm mass between the rheometer plates.^23^ As a consequence, the measured viscoelastic properties represent a biofilm material where the macroscopic structure was largely destroyed. As the structure was locally retained in small volumes (Fig. 5A), it is unclear which interactions determine the average viscoelastic properties of the homogenized biofilm material. In contrast, the original morphology and internal structure is conserved when measuring macrocolony biofilms with microindentation. While this approach accounts for the contribution of biofilm architecture to the mechanical properties measured, the many wrinkles present on biofilms containing pEtN-cellulose, especially when combined with curli, may contribute to the large distribution of values for the different mechanical parameters.^16^ To reduce variations caused by the three-dimensional morphology, we indented biofilms in their central region, where wrinkles are less pronounced and less likely to affect the indentation curves. To mitigate the possible influence of biofilm morphology, we also increased the number of statistical repeats to 10 measurements per biofilm. The positions of these measurements were separated by at least 250 µm to avoid any cross-influence between measurement points.

These technical difficulties related to the complexity of the biofilm material and its morphology add to the variability inherent to biofilm research that has been reported for mechanical studies.^33^ Such variability can be the consequence of subtle changes in experimental conditions, e.g., the atmospheric conditions in the laboratory that can, in turn, affect the (micro)conditions of growth for the bacteria in the agar plate. Indeed, despite following the same protocols as in the literature, the differences in biofilm morphologies, obtained for the different strains, were not as marked as those reported.^35,40^ Moreover, we noticed plate-to-plate variations that we could not attribute to variations in the agar mechanical properties (Fig. S6). Yet, we accounted for possible influences by growing the different *E. coli* strains in each plate, except for the experiment presented in figure 5, where several macrocolony biofilms from one strain were grown in the same plate to compare the two characterization methods. Considering the wide distribution of results, the trends observed in the independent experiments where microindentation and shear-rheology were performed on biofilms from different plates (Fig. 5A), were notably coherent and we are thus confident that the observed differences in mechanical parameters reflect the properties of the biofilms. While no significant influence of homogenization was measured on the reduced indentation elastic moduli of biofilms from different strains (Fig. 5B), we found slightly different trends between strains when comparing the shear moduli obtained by shear-rheology (Fig. 2C, D and Fig.5B) and the reduced elastic moduli obtained by microindentation (Fig. 3C and Fig. 4B). This observation could indicate a major interplay between the loading geometry and biofilm structure during the mechanical test, especially for the samples with no ECM. Differential changes of biofilm adhesion strength observed upon homogenization, as a function of ECM composition (Fig. 5D) may also explain the differences observed while comparing microindentation and shear-rheology. Indeed, while adhesion is not expected to impact the response of the material to compression, it may contribute to the response of the material to shear forces.

Mechanical data obtained at subcellular scales may help to understand biofilm mechanical behavior as a function of their composition. For example, Kreis *et al.*^28^ performed AFM nanoindentation on *E. coli* AR3110 biofilms and identified two populations of ECM with distinct elastic properties (Young’s moduli of ∼1 and 10 kPa, respectively). Taken together with previous observations reported in ^34^, the present results obtained on biofilms from *E. coli* mutants producing either curli or pEtN-cellulose or both types of fibers suggest that the stiffer ECM of *E. coli* AR3110 corresponds to curli fibers while the softer ECM corresponds to pEtN-cellulose. Interestingly, *E. coli* bacteria measured at the subcellular scale are very stiff (∼1 to 10MPa),^28^ although the biofilms grown from *E. coli* deprived of ECM appeared to be the softest of our collection with ∼10kPa (Fig. 3C). In contrast to nanoindentation with AFM, which measures individual bacteria, microindentation on biofilms rather probes their mechanical interactions and how these are mediated by the presence of ECM. As such, a collection of stiff bacteria that are not decorated with ECM fibers is expected to locally rearrange and dissipate energy under minimal compressive load (Fig. 3C and D), while bacteria embedded in curli and/or pEtN-modified cellulose may further transmit forces over larger distances within the biofilm and partially store energy in their ECM. In both cases, the relatively small proportion of force dissipated during the first 205 ms of the holding time indicates that energy dissipation rather occurs *via* bacteria and/or ECM rearrangement than *via* water flow (Fig. 3E).

Our results also suggest that the ability of bacteria to retain or rearrange their organization under mechanical stress depends on the physicochemical properties of their surrounding ECM. For example, cellulose fibers only appear to provide a mechanical advantage to the *E. coli* colony if it is modified with the zwitterionic phosphoethanolamine (pEtN) group. The pEtN-modification not only slightly increases biofilm elastic modulus (Fig. 3C) but also significantly speeds up force relaxation (Fig. 3E) and slightly increases adhesion (Fig. 3F) compared to non-modified cellulose. On the other hand, the presence of curli alone notably increases biofilm rigidity at the expense of a higher plasticity, indicating that the material better resists mechanical stress while it is more prone to irreversible deformation. Finally, our results indicate that biofilms containing both curli and pEtN-cellulose are only stiffer than those containing curli only when the fibers are assembled by the same bacteria (orange data points, Fig. 3C). This clear increase of rigidity suggests that the biofilm matrix is a composite material. It is not observed when curli are combined with non-modified cellulose. Our study thus quantitatively confirms that the natural pEtN-modification has significant mechanical implications at the biofilm level in wild-type strains (AR3110), which is consistent with the observed morphology of macrocolony biofilms (Fig. 1A and 4).^35^ Indeed, wrinkling theory predicts more surface instabilities in stiffer (bio)films, given a similar strain mismatch and adhesion energy at the interface.^41^ The mechanical analysis of biofilms with different combinations of curli and pEtN-cellulose or non-modified cellulose also highlights the importance of when and how the different ECM fibers interact with each other or maybe even co-assemble. When comparing biofilms where curli and pEtN-cellulose are produced by the same bacteria or by different subpopulations, a lower stiffness is observed when the fibers are produced by different bacteria (Fig. 3C). As discussed in earlier work,^42^ bacteria producing only one types of fiber may undergo phase separation during biofilm growth, resulting in regions with different properties. Despite recent work performed with synthetic pEtN-cellulose, the finely tuned interaction between curli and pEtN-cellulose remains poorly understood at the molecular level and is the subject of ongoing work.^43,44^

Lastly, our data suggest that the association of curli and pEtN-cellulose enables *E. coli* to build a tunable composite material that can span a wide range of mechanical properties, potentially using only little variations in composition (i.e. curli to pEtN-cellulose ratio) (Fig. 4). This is consistent with the existence of intracellular levels of regulations, affecting either one or both fibers,^45^ as well as extrinsic compounds targeting specifically one fiber.^46^ While the large spread in our data could be a consequence of the heterogeneity and morphology of the biofilms as discussed earlier,^15,28^ it was also visible in the microindentation experiments performed on homogenized biofilms (Fig. 5B) and in rheology experiments (Fig. 6). The presence of both curli and pEtN-cellulose as building blocks may thus allow *E. coli* to finely adjust the biofilm properties in response to the environment.

Overall, our mechanical comparison of *E. coli* macrocolony biofilms with altered ECM composition quantitatively confirms the key role of curli fibers in biofilm rigidity. It also highlights the crucial contribution of pEtN-cellulose, and especially of the pEtN modification, to the mechanical properties and structural stability of biofilms, especially when associated with curli fibers. While further work at the molecular and fiber levels is needed to understand the interactions between biofilm ECM components, our study provides useful insights into how these interactions impact macrocolony biofilm mechanics. The knowledge presented here is of particular interest for researchers aiming at bridging the gap between ECM production, assembly and hierarchical structure up to macroscopic biofilm materials properties and functions, be it for preventing biofilm formation or engineering them for technical purposes.^47,48^

## Methods

### Bacterial strains and growth

All *E. coli* strains are derived from the strain W3110, which synthesizes amyloid curli protein but no pEtN-cellulose.^16^ Curli amyloid fibers are macromolecular assemblies of csgA (as the major subunit), and csgB (as a minor subunit). The *E. coli* strain AR3110 is a derivative of W3110 with a restored capacity to produce phosphoethanolamine (pEtN)-modified cellulose.^16^ Thus, AR3110 is a highly proficient biofilm-forming strain that produces both amyloid curli protein and pEtN-cellulose as major ECM components (curli+pEtN-cellulose). AP329 (csgBA::kan) is an AR3110 derivative that is deficient in the production of curli while producing pEtN-cellulose.^16^ This strain has a kanamycin resistance cassette associated with the mutation in the structural curli operon (csgBA). AP472 (bcsG::scar, csgBA::kan) is an AR3110 derivative that is deficient in the production of curli and that produces non-modified cellulose (i.e., without the pEtN-modification).^16^ This strain also has a kanamycin resistance cassette associated with the mutation in the structural curli operon (csgBA). AP470 (bcsG::scar) is an AR3110 derivative that produces amyloid curli protein and non-modified cellulose.^16^ AR198 (bcsA::scar, csgB::cm) is also an AR3110 derivative that is deficient in the production of both curli and pEtN-cellulose.^16^ This strain has a chloramphenicol resistance cassette associated with the mutation in the curli structural gene csgB. In this work, we also refer to AR198 as “No ECM” (Fig. 1A). It may still produce ECM components other than curli and pEtN-cellulose, which are usually less abundant (e.g., pga and colanic acid).^49,50^ For the samples termed “co-seeded” (curli:pEtN-cellulose), the strains W3110 and AP329 were mixed in a 1:1 ratio upon seeding. As previously described, OD_600_ was measured for each liquid culture containing one of the strains.^42^ In the case that their OD_600_ was not identical, both cultures were mixed such that the cell density of each strain was identical in the suspension used for seeding. Inoculation on salt-free agar took place immediately after mixing.

Petri dishes (15 cm diameter) were filled with 100 mL of salt-free agar, composed of 1.8 w/v% bacteriological grade agar-agar (Roth, #2266), 1 w/v% tryptone (Roth, #8952) and 0.5 w/v% yeast extract (Roth, #2363). The agar plates were kept in ambient conditions for 48 h before use. Bacteria were initially streaked out on LB agar (Luria/Miller, Roth, #969) and grown overnight at 37 °C. Liquid cultures were subsequently started from a single colony, using liquid LB medium (Roth, #968). Bacteria were again grown overnight at 37 °C with shaking at 250 rpm. Finally, each plate was inoculated with arrays of four or nine drops (5 µL) of bacterial suspension (OD_600_ ∼ 0.5 after 10x dilution). After inoculation, the excess of water evaporated from the drops and bacteria-rich areas of comparable size (diameter of approximately 3 to 4 mm) were visible on the surface. The *E. coli* biofilms were finally grown for 7 days at 28 °C.

### Biofilm area, dry mass, density and water content

After 7 days of growth, 6 out of 9 biofilms per strain were randomly selected and imaged in bright field with a stereomicroscope (AxioZoomV.16, Zeiss, Germany) using the tiling function of the acquisition software (Zen 2.6 Blue edition, Zeiss, Germany). Biofilm size was quantified by calculating their projected area using a custom-made MATLAB code based on thresholding.^29^ The water content of the biofilms was determined by scraping 9 biofilms per condition from the respective agar substrates. Biofilms were placed in plastic weighing boats, and dried in an oven at 60°C for 3 h. Wet and dry masses (m_wet_, m_dry_) were determined before and after drying and used to calculate the water content as W = (m_wet_ - m_dry_)/m_dry_ x 100%w/w.^51^ To calculate biofilm density, the previously calculated areas were used. Assuming the biofilms have a consistent thickness and distribution across the plate, the mean dry mass of the plate (divided by 9 to approximate the unit mass of each biofilm) was divided by the calculated area of each biofilm to obtain the density, expressed in mg.cm^-2^. All procedures were carried out in four independent experiments.

### Oscillatory shear-rheology of biofilms

Biofilms were either measured directly after growth or stored in the fridge for less than 48 h. For storage at 4 °C, the Petri dishes were sealed with parafilm to prevent evaporation. Depending on the strain, two or three biofilms corresponding to a mass of ∼90 mg were harvested from the agar surface using cell scrapers, transferred into a Petri dish and homogenized by stirring for 30 s.

The rheometer used was a stress-controlled shear rheometer (MCR301, Anton Paar GmbH, Ostfildern, Germany). The bottom plate was equipped with Peltier thermoelectric cooling and the temperature was held constant at 21 °C for every experiment. Once the sample was transferred onto the bottom plate, a channel around the plate was filled with water. A hood was lowered down onto the plate to obtain a closed measurement chamber and maintain a high humidity level. A parallel plate geometry with a top plate of 12 mm diameter was used for every experiment. The gap height between the plates was set to 250 µm. Using oscillatory shear measurements, an amplitude sweep was performed to determine the linear viscoelastic range (LVE) and to extract the plateau values of the storage (G’_0_) and loss (G’’_0_) moduli. The oscillation frequency was 10 rad s^-1^. The strain amplitude was first increased step-wise from 0.01 to 100% with seven points per decade and then decreased again to 0.01%. These cycles were repeated three times per experiment (i.e., six intervals). One complete experiment lasted approximately for 45 min. The data presented in the Results section were extracted from the second interval with increasing strain amplitude. The first cycle was considered as one additional homogenization step of the biofilm. The plateau values G’_0_ and G’’_0_ represent the mean values over a strain range from 0.01 to 0.02%, corresponding to three measurement points.

### Microindentation on biofilms

Freshly grown *E. coli* biofilms were used for the indentation experiments. For each strain (incl. the co-seeded W3110 and AP329 strains), ten measurements were performed in the central region of four biofilms each. The distance between two measurement points was at least 250 μm in x and y directions. The indentation depth was between 10 and 30 µm, i.e., much less than the biofilm thickness (∼100 µm).^52^ A TI-950 Triboindenter (Hysitron Inc.) equipped with an extended stage (XZ-500) and a cono-spherical tip (R = 50 µm) were used to determine the load–displacement curves using the ‘air-indent’ method.^36^ Loading rates were set at 20 μm s^-1^, which translates into loading and unloading times of 10 s. The loading portion of all curves were fitted with a Hertzian contact model over an indentation range of 0 to 10 µm to obtain the reduced elastic modulus E_r_.^29^

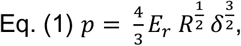

where *p* is the contact force in µN, *E_r_* is the indentation elastic modulus, *R* is the tip radius, and δ is the contact depth. To characterize biofilm relaxation behavior, load-time curves were extracted during the holding period. A local load maximum was consistently observed at 350 ms. It can be attributed to a tip instability, occurring between 250 and 350 ms, when the instrument switches from displacement-controlled indentation to constant displacement for the holding phase. From this curve we calculated the portion of the force relaxation happening before the tip instability as Eq. (2)

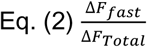

where Δ*F*_*fast*_ represents the fast relaxation reaction occurring between t = 0 – 205 ms (Fig. S3). Δ*F*_*fast*_ was quantified following Eq. (3)

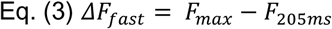

We define *F*_*Total*_ as the total force relaxed after 10 s (endpoint for the holding time), following Eq. (4).

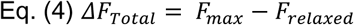

The plasticity index at the holding point (ψ_h_) is defined as Eq. (5) (Fig. S3)

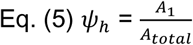

where A_1_ describes the area between the loading and unloading curves during the holding period, and A_total_ describes the area under the loading curve. ψ_h_ spans from 0 to 1, where ψ_h_ = 1 means that the biofilm is fully plastic, and ψ_h_ = 0 means that the biofilm is elastic.

For each biofilm, a subset of curves with an indentation depth between 10 and 25 µm were selected for the analysis of the adhesion force and viscoelastic stress relaxation.

The adhesion strength (*σ*_*Adh*_) is defined as Eq. (6) (Fig. S3).

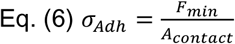

where *F*_*min*_ is the force applied in the detachment point when the tip separates from the biofilm, and *A*_*contact*_ is the contact area between the tip and the biofilm at the largest indentation reached *h*_*max*_ (Fig. S3). *A*_*contact*_ is defined by Eq. (7).

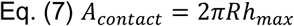

For the analysis of the relaxation curves, plasticity and adhesion strength presented in Fig. 3, only indentation curves that did not exceed 20 µm penetration depth were considered. This was done to avoid any influence of the underlying agar as well as possible deviations from geometrical assumptions on the contact area between the indentation tip and the biofilm.

### Imaging of biofilm cross-sections

Biofilms were grown on agar plates supplemented with the fluorescent dye Direct Red 23 (Pontamine Fast Scarlett 4B) (CAS 3441-14-13, Sigma-Aldrich, Germany), using a final concentration of 0.03 g L^-1^.^53^ Cross-sections of living biofilms were obtained as described previously by Siri et al.^51^ Briefly, liquid plain agar (1.8 % wt) at about 42 °C was slowly poured over each biofilm. The resulting agar−biofilm−agar sandwiches were cut into ∼1 mm-thick slices using a blade. The cross-sections were placed onto glass cover slips. The different cross-sections were observed with a confocal microscope (SP8 FALCON, Leica, Mannheim, Germany), using a 63x oil immersion objective (1.2 NA). Direct Red 23 fluorescence was excited at 552 nm and the emitted photons were collected in the range from 600 to 700 nm. Images were taken and analyzed with the LAS X software (Leica). Four cross-sections were imaged per biofilm.

To image cross-sections of homogenized biofilms, the biofilms were first treated as described in the rheology section. The homogenized biofilms were then transferred into 2 mm wells punctured into a Petri dish of plain agar gel (1.8 % wt) and covered with another layer of agar. The resulting agar−homogenized biofilm−agar sandwiches were cut into ∼1 mm-thick slices using a blade and imaged with a confocal microscope.

### Statistical analysis

To analyze the different parameters obtained from shear-rheology, i.e., plateaus of the storage (G’_0_), loss (G"_0_), and shear moduli (G), yield stress (τ_Y_) and flow stress (τ_F_), the respective median errors were calculated by bootstrapping. From each original data sample X of size 3 (i.e., 3 independent measurements per bacteria strain), 3 elements were chosen with replacement to create a bootstrapped sample Y of size 3. The median m of Y was calculated and stored in the vector M of size 27. The previous steps were repeated for all 27 (= 33) combinations, hence the size of M. The standard deviation of M was taken as the error associated to the median of X.

For each microindentation experiment, N = 3 or 4 biofilms were tested and n = 7 or 10 curves were acquired per biofilm (see figure captions). The mechanical data was plotted and analyzed using the GraphPad Prism 9.3.1 software. Mann-Whitney U tests were used for statistical analysis (p<0.0001, **** | p<0.001, *** | p<0.01, ** | p<0.05, * | ns = non-significant).

## Contributions

Conceptualization: R.Z., A.S., M.S., K.G.B. and C.M.B.

Experimental developments: R.Z., A.S., M.S., S.A.

Data acquisition and analysis: R.Z., A.S., M.S., L.Z., C.S.G.,

Manuscript preparation and writing: M.S., R.Z., A.S., L.Z., C.S.G., C.M.B.

Manuscript reviewing and editing: all co-authors

## Acknowledgements

The authors thank Christine Pilz-Allen for her technical support in the laboratories.

M.S. acknowledges support from the Max Planck Queensland Centre (MPQC) on the Materials Science for Extracellular Matrices. L.Z. acknowledges the research unit “InterDent” and the financial support of the German Research Foundation (Deutsche Forschungsgemeinschaft – DFG, FOR2804). The authors are also grateful for the support of the Cluster of Excellence Matters of Activity. Image Space Material funded by the German Research Foundation (DFG) under Germany’s Excellence Strategy, EXC 2025.

## References

1. Boudarel H, Mathias JDD, Blaysat B, Grediac M, Grédiac M. Towards standardized mechanical characterization of microbial biofilms: analysis and critical review. NPJ Biofilms Microbiomes. 2018;4(1):17. doi:10.1038/s41522-018-0062-5

2. Wolcott RD, Rumbaugh KP, James G, et al. Biofilm maturity studies indicate sharp debridement opens a time-dependent therapeutic window. J Wound Care. 2013;19(8):320–328. doi:10.12968/JOWC.2010.19.8.77709

3. Nerenberg R. The membrane-biofilm reactor (MBfR) as a counter-diffusional biofilm process. Curr Opin Biotechnol. 2016;38:131–136. doi:10.1016/J.COPBIO.2016.01.015

4. Rupp CJ, Fux CA, Stoodley P. Viscoelasticity of Staphylococcus aureus biofilms in response to fluid shear allows resistance to detachment and facilitates rolling migration. Appl Environ Microbiol. 2005;71(4):2175–2178. doi:10.1128/AEM.71.4.2175-2178.2005

5. Even C, Marlière C, Ghigo JM, Allain JM, Marcellan A, Raspaud E. Recent advances in studying single bacteria and biofilm mechanics. Adv Colloid Interface Sci. 2017;247:573–588. doi:10.1016/J.CIS.2017.07.026

6. Gloag ES, Fabbri S, Wozniak DJ, Stoodley P. Biofilm mechanics: Implications in infection and survival. Biofilm. 2020;2:100017. doi:10.1016/J.BIOFLM.2019.100017

7. Wilking JN, Angelini TE, Seminara A, Brenner MP, Weitz DA. Biofilms as complex fluids. MRS Bull. 2011;36(5):385–391. doi:10.1557/mrs.2011.71

8. Tuson HHH, Auer GKK, Renner LDD, et al. Measuring the stiffness of bacterial cells from growth rates in hydrogels of tunable elasticity. Mol Microbiol. 2012;84(5):874–891. doi:10.1111/j.1365-2958.2012.08063.x

9. Bock N, Delbianco M, Eder M, et al. A materials science approach to extracellular matrices. Prog Mater Sci. 2025;149:101391. doi:10.1016/J.PMATSCI.2024.101391

10. Gordon VD, Davis-Fields M, Kovach K, Rodesney CA. Biofilms and mechanics: a review of experimental techniques and findings. J Phys D Appl Phys. 2017;50(22). doi:10.1088/1361-6463/aa6b83

11. Kovach K, Davis-Fields M, Irie Y, et al. Evolutionary adaptations of biofilms infecting cystic fibrosis lungs promote mechanical toughness by adjusting polysaccharide production. npj Biofilms Microbiomes. 2017;3(1):1. doi:10.1038/s41522-016-0007-9

12. Cao H, Habimana O, Safari A, Heffernan R, Dai Y, Casey E. Revealing region-specific biofilm viscoelastic properties by means of a micro-rheological approach. npj Biofilms Microbiomes 2016 21. 2016;2(1):1-7. doi:10.1038/s41522-016-0005-y

13. Siri M, Mangiarotti A, Vázquez-Dávila M, Bidan CM. Curli Amyloid Fibers in Escherichia coli Biofilms: The Influence of Water Availability on their Structure and Functional Properties. Macromol Biosci. 2024;24(2):2300234. doi:10.1002/MABI.202300234

14. Cense AW, Peeters EAG, Gottenbos B, Baaijens FPT, Nuijs AM, van Dongen MEH. Mechanical properties and failure of Streptococcus mutans biofilms, studied using a microindentation device. J Microbiol Methods. 2006;67(3):463–472. doi:10.1016/J.MIMET.2006.04.023

15. Klauck G, Serra DO, Possling A, Hengge R. Spatial organization of different sigma factor activities and c-di-GMP signalling within the three-dimensional landscape of a bacterial biofilm. Open Biol. 2018;8(8). doi:10.1098/rsob.180066

16. Serra DOO, Richter AMM, Hengge R. Cellulose as an architectural element in spatially structured Escherichia coli biofilms. J Bacteriol. 2013;195(24):5540–5554. doi:10.1128/JB.00946-13

17. Serra DO, Hengge R. Cellulose in Bacterial Biofilms. (Cohen E, Merzendorfer H, eds.). Springer; 2019. doi:10.1007/978-3-030-12919-4_8

18. Horvat M, Pannuri A, Romeo T, Dogsa I, Stopar D. Viscoelastic response of Escherichia coli biofilms to genetically altered expression of extracellular matrix components. Soft Matter. 2019;15(25):5042–5051. doi:10.1039/C9SM00297A

19. Rooney LM, Amos WB, Hoskisson PA, McConnell G. Intra-colony channels in E. coli function as a nutrient uptake system. ISME J 2020 1410. 2020;14(10):2461-2473. doi:10.1038/s41396-020-0700-9

20. Wilking JNN, Zaburdaev V, De Volder M, Losick R, Brenner MPP, Weitz DAA. Liquid transport facilitated by channels in Bacillus subtilis biofilms. Proc Natl Acad Sci U S A. 2013;110(3):848–852. doi:10.1073/pnas.1216376110

21. Azulay DN, Spaeker O, Ghrayeb M, et al. Multiscale X-ray study of Bacillus subtilis biofilms reveals interlinked structural hierarchy and elemental heterogeneity. Proc Natl Acad Sci U S A. 2022;119(4):e2118107119. doi:10.1073/PNAS.2118107119

22. Safari A, Tukovic Z, Walter M, Casey E, Ivankovic A. Mechanical properties of a mature biofilm from a wastewater system: from microscale to macroscale level. Biofouling. 2015;31(8):651–664. doi:10.1080/08927014.2015.1075981

23. Lieleg O, Caldara M, Baumgärtel R, Ribbeck K. Mechanical robustness of Pseudomonasaeruginosa biofilms. Soft Matter. 2011;7(7):3307. doi:10.1039/c0sm01467b

24. Kretschmer M, Lieleg O. Chelate chemistry governs ion-specific stiffening of: Bacillus subtilis B-1 and Azotobacter vinelandii biofilms. Biomater Sci. 2020;8(7):1923–1933. doi:10.1039/c9bm01763a

25. Yan J, Moreau A, Khodaparast S, et al. Bacterial Biofilm Material Properties Enable Removal and Transfer by Capillary Peeling. Adv Mater. 2018;30(46):e1804153. doi:10.1002/adma.201804153

26. Tallawi M, Opitz M, Lieleg O. Modulation of the mechanical properties of bacterial biofilms in response to environmental challenges. Biomater Sci. 2017;5(5):887–900. doi:10.1039/C6BM00832A

27. Kretschmer M, Schüßler CA, Lieleg O. Biofilm Adhesion to Surfaces is Modulated by Biofilm Wettability and Stiffness. Adv Mater Interfaces. 2021;8(5). doi:10.1002/admi.202001658

28. Kreis CTT, Sullan RMAMA. Interfacial nanomechanical heterogeneity of the E. coli biofilm matrix. Nanoscale. 2020;12(32):16819–16830. doi:10.1039/D0NR03646C

29. Ziege R, Tsirigoni AM, Large B, et al. Adaptation of Escherichia coli Biofilm Growth, Morphology, and Mechanical Properties to Substrate Water Content. ACS Biomater Sci Eng. 2021;7(11):5315–5325. doi:10.1021/ACSBIOMATERIALS.1C00927

30. Peterson BW, van der Mei HC, Sjollema J, Busscher HJ, Sharma PK. A distinguishable role of eDNA in the viscoelastic relaxation of biofilms. MBio. 2013;4(5). doi:10.1128/MBIO.00497-13

31. Rogers SS, van der Walle C, Waigh TA. Microrheology of Bacterial Biofilms In Vitro: Staphylococcus aureus and Pseudomonas aeruginosa. Langmuir. 2008;24(23):13549–13555. doi:10.1021/la802442d

32. Galy O, Latour-Lambert P, Zrelli K, Ghigo JM, Beloin C, Henry N. Mapping of Bacterial Biofilm Local Mechanics by Magnetic Microparticle Actuation. Biophys J. 2012;103(6):1400–1408. doi:10.1016/j.bpj.2012.07.001

33. Qi L, Christopher GF. Rheological variability of Pseudomonas aeruginosa biofilms. Rheol Acta. 2021;60(4):219–230. doi:10.1007/S00397-021-01260-W

34. Serra DO, Richter AM, Hengge R. Cellulose as an architectural element in spatially structured escherichia coli biofilms. J Bacteriol. 2013;195(24):5540–5554. doi:10.1128/JB.00946-13

35. Thongsomboon W, Serra DO, Possling A, Hadjineophytou C, Hengge R, Cegelski L. Phosphoethanolamine cellulose: A naturally produced chemically modified cellulose. Science (80- ). 2018;359(6373):334-338. doi:10.1126/science.aao4096

36. Amini S, Chen X, Chua JQI, Tee JS, Nijhuis CA, Miserez A. Interplay between Interfacial Energy, Contact Mechanics, and Capillary Forces in EGaIn Droplets. ACS Appl Mater Interfaces. 2022;14(24):28074–28084. doi:10.1021/ACSAMI.2C04043/

37. Peterson BW, Busscher HJ, Sharma PK, Van Der Mei HC. Visualization of Microbiological Processes Underlying Stress Relaxation in Pseudomonas aeruginosa Biofilms. Microsc Microanal. 2014;20(3):912–915. doi:10.1017/S1431927614000361

38. Jana S, Charlton SGV, Eland LE, et al. Nonlinear rheological characteristics of single species bacterial biofilms. npj Biofilms Microbiomes. 2020;6(1):1–11. doi:10.1038/S41522-020-0126-1/

39. Geisel S, Secchi E, Vermant J. Experimental challenges in determining the rheological properties of bacterial biofilms. Interface Focus. 2022;12(6). doi:10.1098/RSFS.2022.0032

40. Serra DO, Richter AM, Klauck G, Mika F, Hengge R. Microanatomy at Cellular Resolution and Spatial Order of Physiological Differentiation in a Bacterial Biofilm. MBio. 2013;4(2):1–12. doi:10.1128/mBio.00103-13

41. Wang Q, Zhao X. A three-dimensional phase diagram of growth-induced surface instabilities. Sci Rep. 2015;5:8887. doi:10.1038/srep08887

42. Sarlet A, Ruffine V, Blank KG, Bidan CM. Influence of Metal Cations on the Viscoelastic Properties of Escherichia coli Biofilms. ACS Omega. 2023;8(5):4667–4676. doi:10.1021/ACSOMEGA.2C06438

43. Tyrikos-Ergas T, Gim S, Huang JY, et al. Synthetic phosphoethanolamine-modified oligosaccharides reveal the importance of glycan length and substitution in biofilm-inspired assemblies. Nat Commun 2022 131. 2022;13(1):1-8. doi:10.1038/s41467-022-31633-5

44. Adams CE, Spicer SK, Gaddy JA, Townsend SD. Synthesis of a Phosphoethanolamine Cellulose Mimetic and Evaluation of Its Unanticipated Biofilm Modulating Properties. ACS Infect Dis. Published online 2024. doi:10.1021/ACSINFECDIS.4C00267

45. Hengge R. Linking bacterial growth, survival, and multicellularity – small signaling molecules as triggers and drivers. Curr Opin Microbiol. 2020;55:57–66. doi:10.1016/J.MIB.2020.02.007

46. Cordisco E, Zanor MI, Moreno DM, Serra DO. Selective inhibition of the amyloid matrix of Escherichia coli biofilms by a bifunctional microbial metabolite. npj Biofilms Microbiomes 2023 91. 2023;9(1):1-14. doi:10.1038/s41522-023-00449-6

47. Wells M, Schneider R, Bhattarai B, et al. Perspective: The viscoelastic properties of biofilm infections and mechanical interactions with phagocytic immune cells. Front Cell Infect Microbiol. 2023;13:1102199. doi:10.3389/FCIMB.2023.1102199

48. An B, Wang Y, Huang Y, et al. Engineered Living Materials For Sustainability. Chem Rev. 2023;123(5):2349–2419. doi:10.1021/ACS.CHEMREV.2C00512/

49. Wang X, Preston JF, Romeo T. The pgaABCD Locus of Escherichia coli Promotes the Synthesis of a Polysaccharide Adhesin Required for Biofilm Formation. J Bacteriol. 2004;186(9):2724–2734. doi:10.1128/JB.186.9.2724-2734.2004

50. Prigent-Combaret C, Prensier G, Le Thi TT, Vidal O, Lejeune P, Dorel C. Developmental pathway for biofilm formation in curli-producing Escherichia coli strains: role of flagella, curli and colanic acid. Environ Microbiol. 2000;2(4):450–464. doi:10.1046/J.1462-2920.2000.00128.X

51. Siri M, Vázquez-Dávila M, Sotelo Guzman C, Bidan CM. Nutrient availability influences E. coli biofilm properties and the structure of purified curli amyloid fibers. npj Biofilms Microbiomes 2024 101. 2024;10(1):1-12. doi:10.1038/s41522-024-00619-0

52. Serra DOO, Klauck G, Hengge R. Vertical stratification of matrix production is essential for physical integrity and architecture of macrocolony biofilms of Escherichia coli. Env Microbiol. 2015;17(12):5073–5088. doi:10.1111/1462-2920.12991

53. Serra DO, Hengge R. A c-di-GMP-Based Switch Controls Local Heterogeneity of Extracellular Matrix Synthesis which Is Crucial for Integrity and Morphogenesis of Escherichia coli Macrocolony Biofilms. J Mol Biol. 2019;431(23):4775–4793. doi:10.1016/J.JMB.2019.04.001

