## Supplement Information file for "Mechanical comparison of *Escherichia coli* biofilms with altered matrix composition: a study combining shear-rheology and microindentation"

### Supplementary Information

| | |
|---|---|
| Biofilm area: Statistical significance | Table S1 |
| Biofilm dry mass: Statistical significance | Table S2 |
| Biofilm density: Statistical significance | Table S3 |
| Biofilm water contents and water activities | Figure S1 |
| Shear modulus $G_0$ calculated from $G'_0$ and $G''_0$ | Figure S2 |
| Microindentation curves | Figure S3 |
| Biofilm mechanical properties: Statistical differences | Tables S4-S7 |
| Quantification of biofilm wrinkling | Figure S4 |
| | Table S8 |
| Influence of a homogenization step on biofilm mechanical properties | Figure S5 |
| | Table S9 |
| Comparative plot of reduced elastic moduli of biofilms vs agar plates | Figure S6 |

**Table S1.** Statistical significance for Fig. 1B; Biofilm area.

| Area (mm²) | AR198 | AP472 | AP329 | W3110 | AP470 | AR3110 | 50:50 |
|---|---|---|---|---|---|---|---|
| AR198 | | * | ns | ns | ns | * | ns |
| AP472 | * | | ns | * | ** | *** | ns |
| AP329 | ns | ns | | ns | ** | *** | ns |
| W3110 | ns | * | ns | | ns | * | ns |
| AP470 | ns | ** | ** | ns | | ns | ns |
| AR3110 | * | *** | *** | * | ns | | * |
| 50:50 | ns | ns | ns | ns | ns | * | |

**Table S2.** Statistical significance for Fig. 1C; Biofilm dry mass.

| Dry mass (mg) | AR198 | AP472 | AP329 | W3110 | AP470 | AR3110 | 50:50 |
|---|---|---|---|---|---|---|---|
| AR198 | | ns | ns | ns | ns | ns | ns |
| AP472 | ns | | ns | ns | ns | ns | ns |
| AP329 | ns | ns | | ns | ns | ns | ns |
| W3110 | ns | ns | ns | | ns | ns | ns |
| AP470 | ns | ns | ns | ns | | ns | ns |
| AR3110 | ns | ns | ns | ns | ns | | ns |
| 50:50 | ns | ns | ns | ns | ns | ns | |

**Table S3.** Statistical significance for Fig. 1D; Biofilm density.

| Density (mg/cm$^{-2}$) | AR198 | AP472 | AP329 | W3110 | AP470 | AR3110 | 50:50 |
|---|---|---|---|---|---|---|---|
| AR198 | | ns | *** | * | * | ns | ns |
| AP472 | ns | | ** | ns | * | ns | ns |
| AP329 | *** | ** | | **** | **** | ** | *** |
| W3110 | * | ns | **** | | ns | ns | ns |
| AP470 | * | * | **** | ns | | ns | * |
| AR3110 | ns | ns | ** | ns | ns | | ns |
| 50:50 | ns | ns | *** | ns | * | ns | |

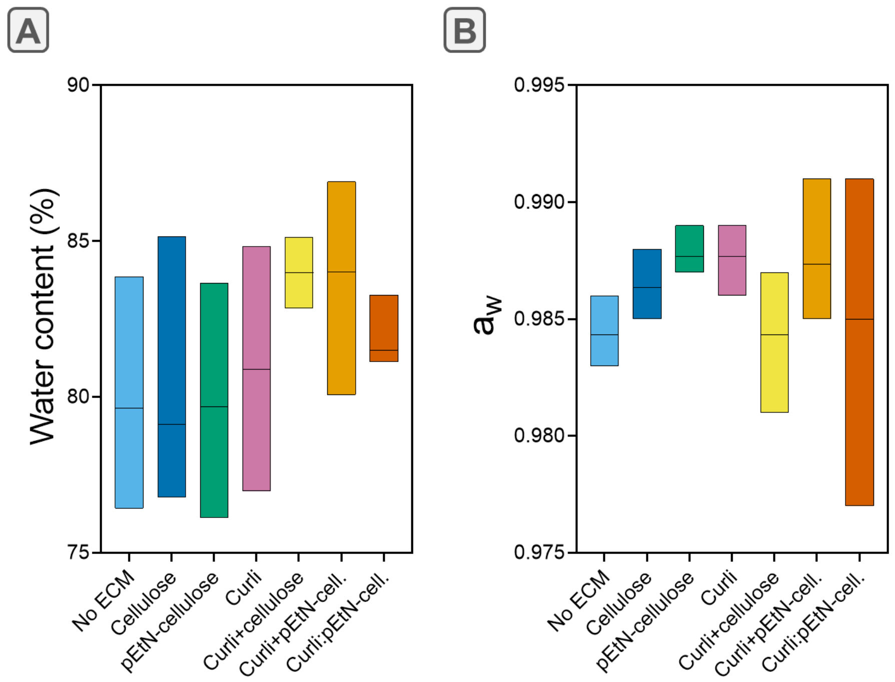


**Figure S1.** Biofilm water contents (A) and water activities (B). Differences were not statistically significant.

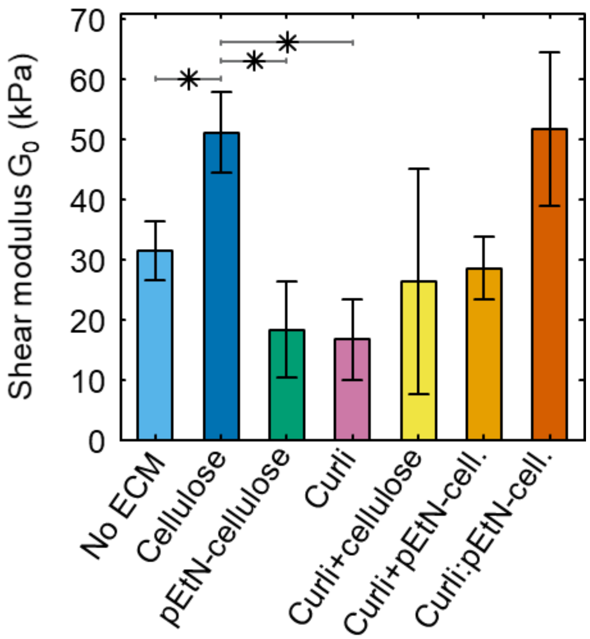


**Figure S2.** Shear Moduli $G_0$ calculated from $G'_0$ and $G''_0$ ($G_0 = \sqrt{{G_0}'^2 + {G_0}''^2}$)

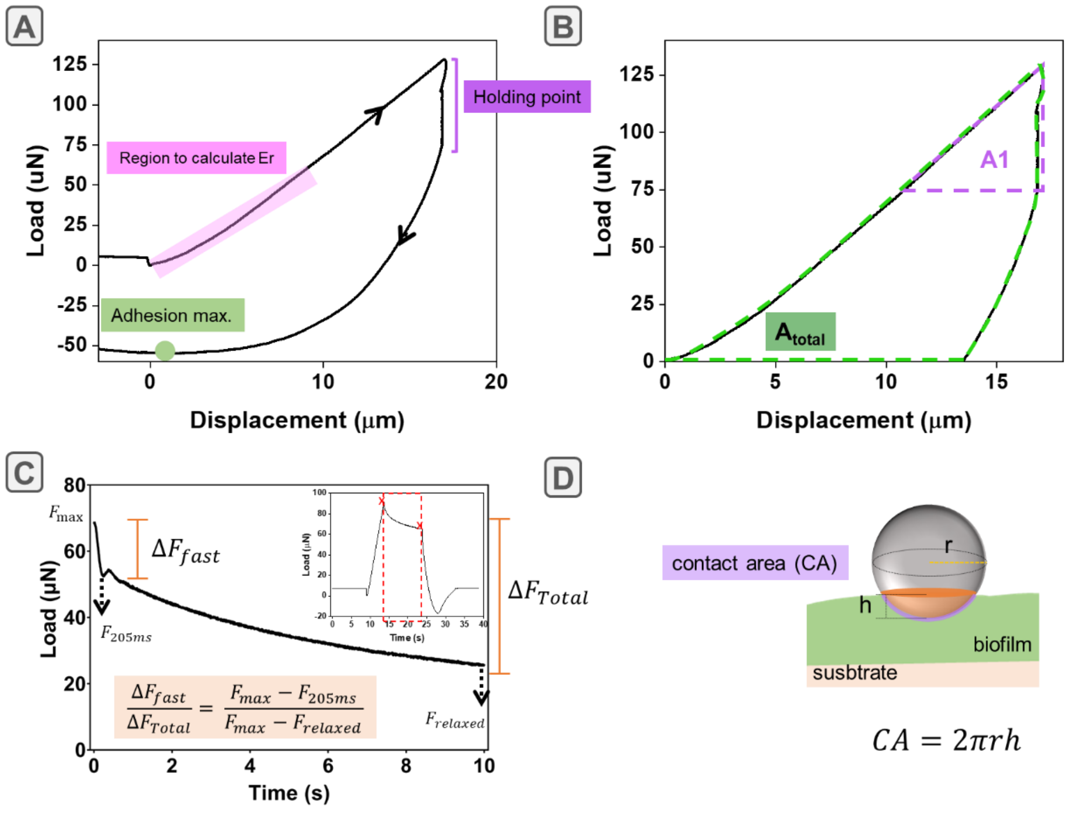


**Figure S3.** Microindentation of biofilms. (A) Representative load-displacement curve of a biofilm. A holding time of 10 s was set between loading and retraction of the tip. (B) Areas used for calculating the plasticity during the holding time: A1 is defined by the quasi-triangle formed by the loading curve and the load relaxation during the holding time and $A_{total}$ is the area under the loading and detachment curves. (C) Analysis of biofilm relaxation behavior during the holding time. (D) Details of the contact area calculation used to determine the adhesion strength.

**Table S4.** Statistical significance for Fig. 3C; Biofilm reduced modulus.

| $E_r$ [kPa] | AR198 | AP472 | AP329 | W3110 | AP470 | AR3110 | 50:50 |
|---|---|---|---|---|---|---|---|
| AR198 | | ns | **** | **** | **** | **** | **** |
| AP472 | ns | | **** | **** | **** | **** | **** |
| AP329 | **** | **** | | **** | **** | **** | ns |
| W3110 | **** | **** | **** | | ns | **** | ns |
| AP470 | **** | **** | **** | ns | | **** | ns |
| AR3110 | **** | **** | **** | **** | **** | | **** |
| 50:50 | **** | **** | ns | ns | ns | **** | |

**Table S5.** Statistical significance for Fig. 3E; ΔFN.

| $F_{fast}/F_{total}$ | AR198 | AP472 | AP329 | W3110 | AP470 | AR3110 | 50:50 |
|---|---|---|---|---|---|---|---|
| AR198 | | *** | **** | **** | **** | ns | * |
| AP472 | *** | | **** | ns | ns | **** | ns |
| AP329 | **** | **** | | **** | ** | **** | ** |
| W3110 | **** | ns | **** | | ns | **** | ns |
| AP470 | **** | ns | ** | ns | | **** | ns |
| AR3110 | ns | **** | **** | **** | **** | | ** |
| 50:50 | * | ns | ** | ns | ** | ** | |

**Table S6.** Statistical significance for Fig. 3F; Plasticity at holding time.

| $(\psi_h)$ | AR198 | AP472 | AP329 | W3110 | AP470 | AR3110 | 50:50 |
|---|---|---|---|---|---|---|---|
| AR198 | | ns | ** | ns | ns | ns | ns |
| AP472 | ns | | * | ns | ns | ns | ns |
| AP329 | ** | * | | *** | *** | *** | ns |
| W3110 | ns | ns | *** | | ns | ns | ns |
| AP470 | ns | ns | *** | ns | | ns | ns |
| AR3110 | ns | ns | *** | ns | ns | | ns |
| 50:50 | ns | ns | ns | ns | ns | ns | |

**Table S7.** Statistical significance for Fig. 3G; Adhesion strength.

| $\sigma_{Adh}$ (kPa) | AR198 | AP472 | AP329 | W3110 | AP470 | AR3110 | 50:50 |
|---|---|---|---|---|---|---|---|
| AR198 | | ns | ns | ns | *** | ns | ns |
| AP472 | ns | | ns | ns | **** | ns | * |
| AP329 | ns | ns | | ns | * | ns | ns |
| W3110 | ns | ns | ns | | * | ns | ns |
| AP470 | *** | **** | * | * | | ** | * |
| AR3110 | ns | ns | ns | ns | ** | | ns |
| 50:50 | ns | * | ns | ns | * | ns | |

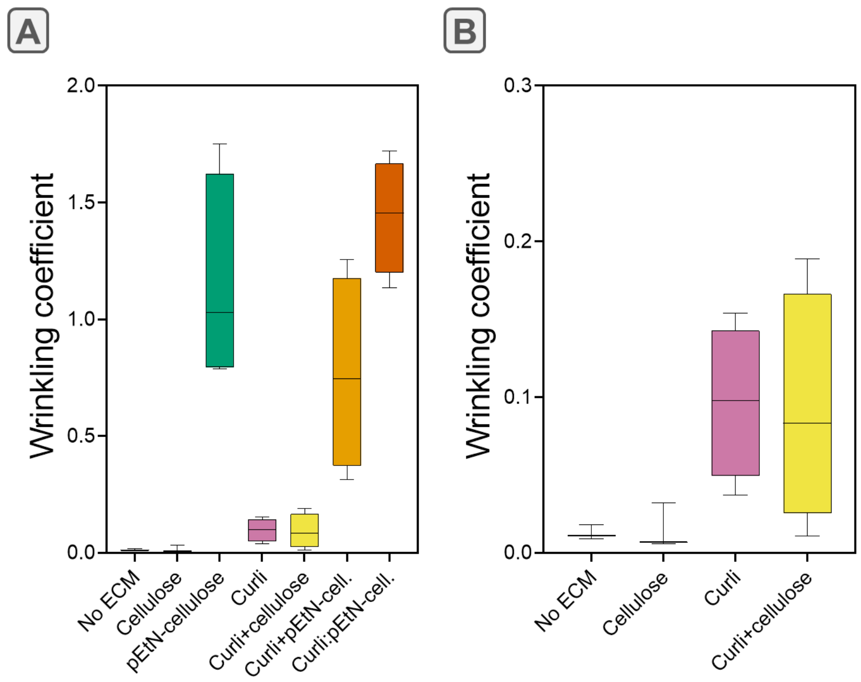


**Figure S4.** (A) Wrinkling quantification from Figure 4; (B) Zoom of A between 0 and 0.3.

**Table S8.** Statistical significance of the wrinkling coefficient in Fig. S4.

| Wrinkling coefficient | AR198 | AP472 | AP329 | W3110 | AP470 | AR3110 | 50:50 |
|---|---|---|---|---|---|---|---|
| AR198 | | ns | *** | ns | ns | * | **** |
| AP472 | ns | | *** | ns | ns | * | **** |
| AP329 | *** | *** | | *** | *** | ns | **** |
| W3110 | ns | ns | *** | | ns | * | **** |
| AP470 | ns | ns | *** | ns | | * | **** |
| AR3110 | * | * | ns | * | * | | * |
| 50:50 | **** | **** | ns | **** | **** | * | |

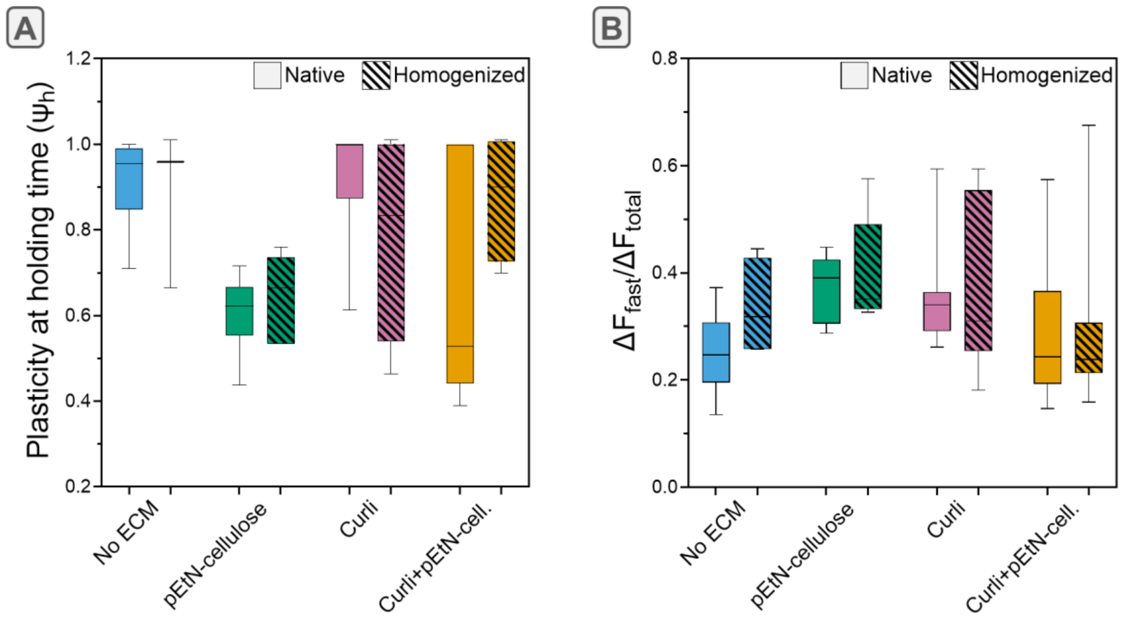


**Figure S5.** Influence of a homogenization step (mixing) on the plasticity (A) and the relaxation behaviour (B) of the biofilm.

**Table S9.** Statistical significance of the comparison between native vs. homogenized biofilms, shown in Fig. 5B, C and D.

| | Reduced modulus, Er (kPa) | Plasticity$_h$ ($\psi_h$) | $\Delta F_{fast}/\Delta F_{total}$ | Adhesion strength (kPa) |
|---|---|---|---|---|
| AR198 Nat vs. Hom. | ns | ns | ns | ns |
| AP329 Nat. vs Hom. | ns | ns | ns | *** |
| AR3110 Nat. vs Hom. | ns | ns | ns | ns |
| W3110 Nat. vs Hom. | ns | ns | * | * |

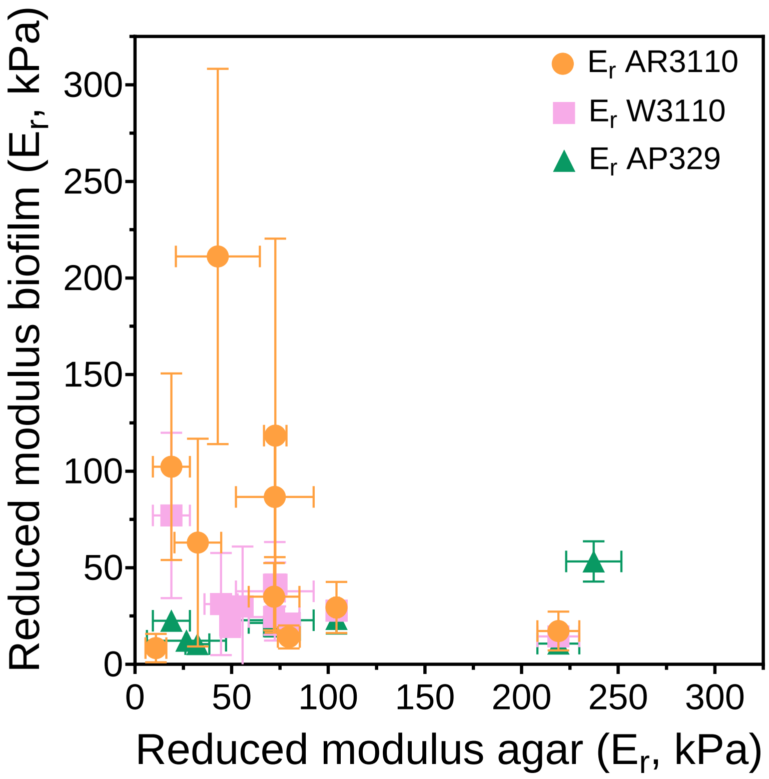


**Figure S6.** Comparative plot of the reduced elastic moduli of biofilms vs. agar plates.
